# FoldAbS: Repurposing the Protein Folding Model as a Foundation Encoder for Antibody Screening

**DOI:** 10.64898/2026.11.03.763899

**Authors:** Jun Wu, Fandi Wu, Xinyuan Zhu, Dawei Huang, Kaiwen Cheng, Jianzhu Ma, Fuli Feng, Jianhua Yao

## Abstract

Therapeutic antibody screening requires prioritizing antibody candidates across specificity, affinity, and developability. Current pipelines typically silo these objectives and underuse structural information that governs antibody–antigen recognition. To address these inefficacies, we introduce **FoldAbS**, a unified supervised screening framework that reconceptualizes protein folding models as frozen foundation encoders, exploiting their internal representations to capture rich sequence, geometric, and interaction dynamics well beyond basic structure prediction. By coupling the folding model with lightweight, task-specific heads, FoldAbS seamlessly evaluates all three therapeutic criteria within a single architecture. Instantiating this framework with the open-source Protenix model (**FoldAbS-Px**) yields consistent improvements over baseline approaches, achieving significant performance gains of up to 13.1%, 9.2%, and 6.2% in specificity, affinity, and developability, respectively. Collectively, these results reposition protein folding models from structure predictors and confidence scorers into reusable foundation encoders for comprehensive and supervised antibody screening. Our code is available at https://github.com/nohandsomewujun/FoldAbS.

## 1 Introduction

Antibody screening (AbS) aims to prioritize a small set of candidates for downstream validation based on antigen-specific binding, sufficient affinity, and favorable developability properties, including stability, solubility, and manufacturability [Tiller and Tessier, 2015, Rabia et al., 2018, Jain et al., 2017]. As candidate libraries continue to expand, AbS has become increasingly critical given the cost and throughput constraints of experimental assays [Raybould et al., 2019, Jain et al., 2017, Chungyoun and Gray, 2025, Yu et al., 2026, Pacesa et al., 2025]. Because antibody–antigen recognition is intrinsically governed by complex structure [Song et al., 2026, Chungyoun and Gray, 2025], effective AbS mandates the rigorous integration of structural information. Existing computational approaches remain limited, either relying on heuristic structural scores or failing to align with screening objectives [Song et al., 2026, He et al., 2024, Agarwal et al., 2024, Luo et al., 2023, Pacesa et al., 2025, Wu et al., 2025, 2024]. Moreover, specificity, affinity, and developability are typically assessed by separate models with heterogeneous inputs and output scales, making screening pipelines computationally cumbersome and difficult to scale to large candidate pools. Overcoming these limitations necessitates a unified framework that comprehensively exploits antibody–antigen structural information while learning task-aligned predictors for specificity, affinity, and developability.

Protein folding models provide a natural substrate for a unified AbS framework, as they inherently encode sequence, geometry, residue packing, intermolecular contacts, and other biophysical regularities [Jumper et al., 2021, Abramson et al., 2024, Zhang et al., 2026, Passaro et al., 2025]. By tightly coupling sequence and structural interaction data, their internal representations are ideally suited for AbS objectives. However, existing AbS methods that leverage folding models still underutilize this rich latent space. Methods based on predicted structures [Song et al., 2026, Li et al., 2025, Zou et al., 2026] force downstream AbS models to relearn interaction features from coordinates, while methods based on confidence scores, such as ipTM or handcrafted interface scores [Pacesa et al., 2025, Team et al., 2025, Yu et al., 2026, Dunbrack Jr, 2025, Wu et al., 2024, Fang et al., 2025] compress nuanced structural uncertainty and interaction information into simplistic ranking values that are not optimized for screening objectives. To fully harness the biophysical priors embedded within protein folding models, their internal representations must be directly leveraged as the primary substrate for supervised AbS.

Adapting these internal representations for AbS is constrained by two fundamental challenges. First, the three screening objectives rely on divergent signals—interaction geometry for specificity and affinity, versus antibody-intrinsic biophysical patterns for developability. Second, specificity prediction necessitates supervised binder/non-binder labels despite a severe scarcity of experimentally validated negative pairs. To address these challenges, we propose **FoldAbS**, a unified framework for supervised AbS anchored by a frozen folding model as its foundational encoder. To accommodate the distinct signal requirements of each screening objective, FoldAbS integrates lightweight, taskspecific learning heads. We enable supervised specificity learning by deriving robust pseudo-negative training signals from observed binders using complementary generation strategies. We instantiate the framework as **FoldAbS-Px** with Protenix [Zhang et al., 2026] and conduct extensive evaluation on six benchmarks, establishing a new state-of-the-art with significant performance improvements across specificity, affinity, and developability. In summary, **our key contributions are as follows**:

- We propose **FoldAbS**, a supervised AbS framework that unifies specificity, affinity, and developability prediction by adapting a protein folding model into a task-aware foundation encoder.
- We present **FoldAbS-Px**, an instantiation of this framework, and design complementary generation strategies to derive robust pseudo-negative training signals for supervised specificity learning.
- We validate the rationality and effectiveness of our proposal via extensive experiments on six benchmarks, pushing the paradigm shift of AbS to unified folding architectures.

## 2 Related Work

### Specificity and affinity prediction

Specificity and affinity prediction assess whether an antibody binds a target antigen and how strongly it binds, forming the core interaction-level criteria for antibody screening. Existing approaches range from sequence-based models, especially protein language models, such as A2binder [He et al., 2024], S3AI [Liu et al., 2024], Ophiuchus-Ab [Zhu et al., 2026], and AlphaBind [Agarwal et al., 2024], to structure-aware methods that exploit antibody– antigen complexes, interface features, geometric descriptors, or folding-model confidence scores as proxy ranking signals, including InterAb [Song et al., 2026] and recent confidence-based screening workflows [Dunbrack Jr, 2025, Pacesa et al., 2025, Yu et al., 2026]. Sequence-based models typically extract antibody and antigen embeddings from pretrained encoders and fine-tune task-specific heads for specificity or affinity estimation [Liu et al., 2026, He et al., 2024], making them scalable but limited in explicit access to molecular recognition geometry. Structure-aware methods better capture intermolecular contacts, but often treat structures or confidence scores as fixed inputs rather than adapting the transferable internal representations learned by folding models.

### Antibody developability prediction

Antibody developability prediction focuses on biophysical and translational properties such as stability, aggregation, expression, and immunogenicity. Existing approaches are typically formulated as task-specific supervised models using sequence-derived descriptors, physicochemical features, or pretrained protein and antibody language model embeddings [Ruffolo et al., 2021, Lin et al., 2023]. Recent foundation-model-based work, such as OphiuchusAb [Zhu et al., 2026], suggests that pretrained antibody representations can also support developability prediction. However, large-scale benchmarking in FLAb2 [Chungyoun and Gray, 2025] shows that developability remains difficult to predict consistently across datasets and properties, with no single model performing robustly across all settings.

### Protein folding models in antibody screening

Recent protein folding models, represented by AlphaFold 3 [Abramson et al., 2024] and related open-source models [Passaro et al., 2025, Zhang et al., 2026, team et al., 2024, The OpenFold3 Team, 2025, Qiao et al., 2026], have greatly expanded access to structure prediction for biomolecular complexes. In antibody screening, these models are commonly used in two ways. First, predicted antibody–antigen complex structures serve as structural inputs for downstream scoring models. Second, native confidence or interface-level scores, such as ipTM and ipSAE [Dunbrack Jr, 2025], are directly used to rank candidate binders or filter designed sequences [Pacesa et al., 2025, Team et al., 2025, Yu et al., 2026]. These applications have made folding models important components of structure-aware screening pipelines. AF3-like folding models have also been adapted for antibody sequence–structure co-design [Yang et al., 2026]. However, most existing uses still treat them as structure generators or confidence-based scoring proxies, leaving their potential as reusable foundation encoders for supervised antibody screening less explored.

## 3 Methodology

As shown in Figure 1, FoldAbS-Px uses Protenix-v1 [Zhang et al., 2026] as a fixed structural encoder and trains lightweight heads for three antibody screening tasks: specificity, affinity, and developability prediction.

**Figure 1:**
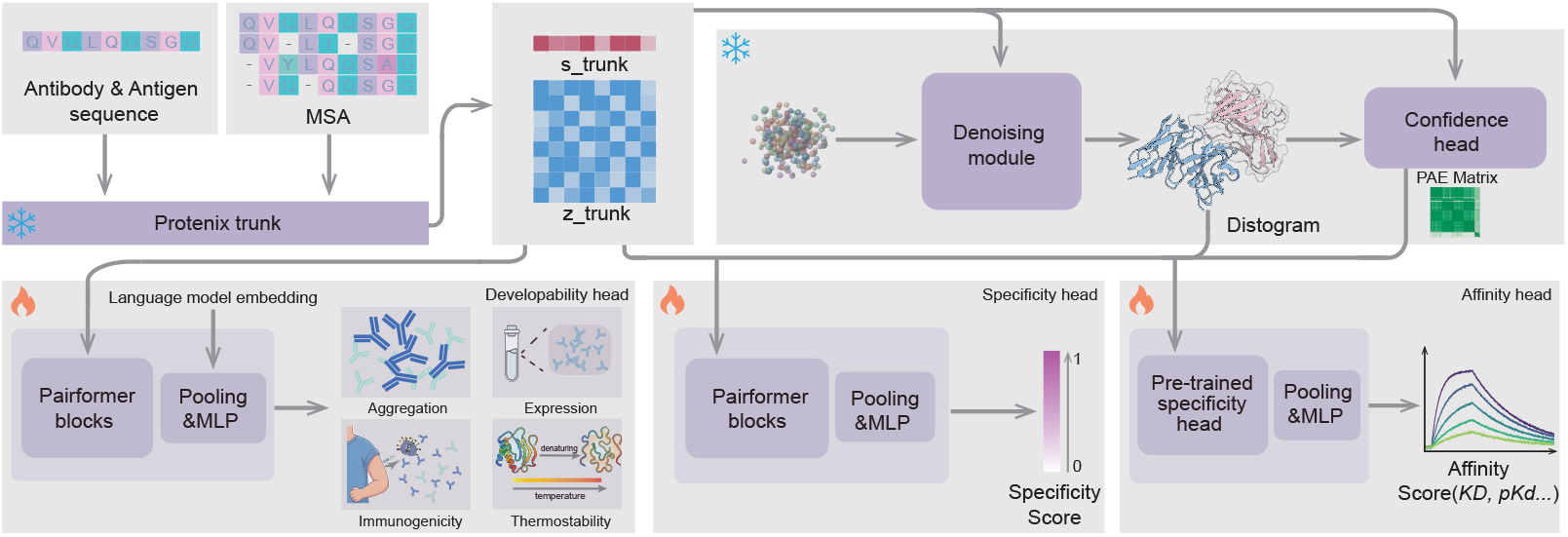
Overview of FoldAbS-Px. FoldAbS-Px uses a frozen Protenix model as a shared foundation encoder for antibody specificity, affinity, and developability prediction. Given antibody and antigen inputs, Protenix generates trunk representations and structural outputs, including predicted complex structures, distograms, and confidence estimates. Protenix-derived representations are used by taskspecific heads for binder discrimination, developability prediction over aggregation, expression, immunogenicity, and thermostability, and affinity regression, where the affinity head is initialized from the pretrained specificity head to reuse interaction-level representations.

### 3.1 Overview of FoldAbS-Px

Let *x*_ab_denote an antibody input and *x*_ag_ denote an antigen input. Each input consists of an amino-acid sequence and its associated MSA features:

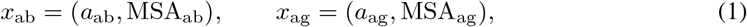

where *a* denotes the amino-acid sequence and MSA denotes precomputed multiple-sequencealignment features used by the folding models. Details of MSA construction are provided in Appendix B.1.

Specificity and affinity are defined as antibody–antigen pair-level tasks with input *x* = (*x*_ab_, *x*_ag_), whereas developability is defined as an antibody-level task with input *x* = *x*_ab_. Specificity prediction is formulated as binary classification between cognate binder pairs and non-binding or non-cognate pairs. Affinity prediction estimates *pK*_*d*_ for antibody–antigen pairs, and developability prediction estimates experimentally measured antibody biophysical readouts, such as immunogenicity (%ADA) and expression level (mg/L).

For compact notation, FoldAbS-Px uses a task-specific feature subset ℛ_*t*_(*x*) and a downstream prediction head:

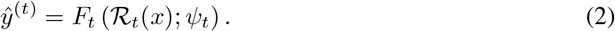

Here, *F*_*t*_ denotes the corresponding task head, with *F*_*t*_ *∈ {F*_spec_, *F*_aff_, 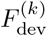}; for developability, *k* indexes the property-specific predictor. The Protenix encoder is kept fixed, and only the downstream parameters *ψ*_*t*_ are optimized.

### 3.2 Protenix-Derived Representations

Let *P*_*θ*_ denote Protenix, whose parameters *θ* are fixed during training. For an input *x*, Protenix produces token-level, pair-level, and structure-derived outputs:

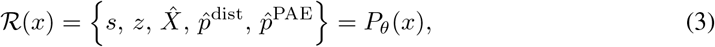

where 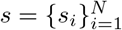 and 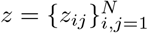 are the single and pair representations from the Protenix trunk; 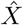 is the predicted atomic structure from the denoising module; 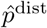 is the distogram distribution produced by the Protenix distogram head; and 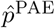 is the predicted aligned error distribution produced by the confidence head. Here, *N* denotes the number of amino-acid residues in the input. A geometry-derived distance map is further computed from the predicted coordinates:

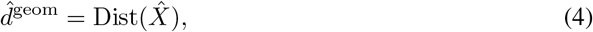

where Dist(*·*) computes pairwise distances from the predicted structure. 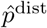 denotes the Protenix distogram output, whereas 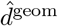 denotes the distance map derived from 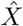.

Specificity and affinity use the complex-level features {*s, z*, 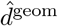, 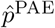} from antibody–antigen inputs, whereas developability uses {*s, z*, 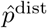} from antibody-only inputs. In FoldAbS-Px, Protenix is kept frozen, and only the downstream heads are trained, preserving the pretrained folding prior while avoiding perturbation of structure-generating representations under limited task-specific supervision.

### 3.3 Task-Specific Prediction Heads

The abstract head *F*_*t*_ introduced above is instantiated separately for specificity, affinity, and each developability property as follows.

#### Specificity head

Since specificity depends on antibody–antigen interaction geometry, FoldAbS-Px uses complex-level features:

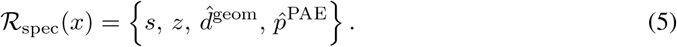

The specificity head maps these features to a binding logit:

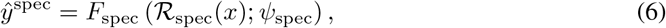

where *ψ*_spec_ denotes the trainable parameters of the specificity head. The binding probability is obtained as *σ*(*ŷ*^spec^), where *σ*(*·*) is the sigmoid function. In contrast to native confidence scores such as ipTM, this head learns a task-aligned decision function for binder discrimination directly from shared structural representations. Empirical evidence and ablations for this design choice are provided in Appendix D.1.

#### Affinity head

Affinity prediction uses the same interaction-level feature set as specificity:

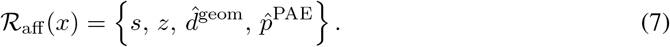

The affinity head *F*_aff_ is initialized from the specificity head because specificity training provides an interaction representation suitable for downstream affinity regression. Architecturally, *F*_aff_ reuses the specificity head up to its penultimate hidden layer and replaces the final classification layer with a regression MLP:

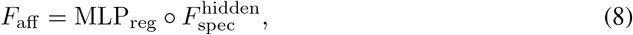

where 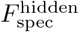 denotes the specificity head before its final classification layer. The affinity head maps the interaction-level features to a continuous binding score:

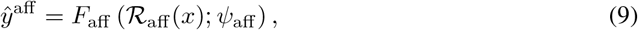

where *ψ*_aff_ denotes the trainable downstream parameters of the affinity head.

The final classification layer is replaced because affinity prediction requires estimating continuous binding strength rather than a binary binder label. During affinity training, the Protenix backbone remained frozen, and all downstream affinity-head parameters *ψ*_aff_ were fine-tuned. This adaptation is important because affinity datasets are composed primarily of binding pairs: a frozen specificity head tends to map these examples to similarly high-confidence binder representations, whereas affinity prediction requires resolving finer differences in binding strength among binders. Fine-tuning the inherited specificity module therefore adapts the interaction embedding from binary binder discrimination to continuous affinity regression. The empirical evidence for this design choice is summarized in Appendix D.2.

#### Developability head

For developability prediction, Protenix is applied to the antibody input *x*_ab_. Since developability is primarily governed by antibody-intrinsic biophysical properties, the developability head uses the antibody-level features

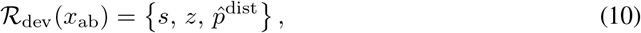

where *s, z*, and 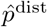 are the Protenix outputs for the antibody input. These design choices are ablated in Appendix D.3.

For the *k*-th developability property, the prediction is

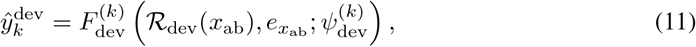

where 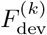 is the predictor for the *k*-th property, 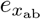 denotes optional antibody sequence embeddings from the language model, and 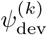 denotes its trainable parameters. Implementation details are provided in Appendix A.2.

### 3.4 Model Training

#### Pseudo-negative construction for specificity supervision

Specificity prediction requires binderversus-non-binder supervision, but large-scale antibody–antigen datasets are dominated by observed binding pairs and contain few experimentally validated non-binders. Let *D*^+^ denote the observed positive set with *N*_+_ antibody–antigen pairs:

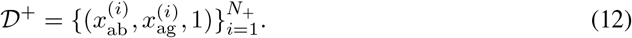

The specificity training set was defined as

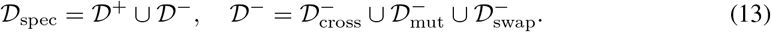

The three pseudo-negative components were constructed as

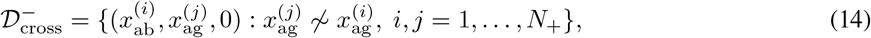

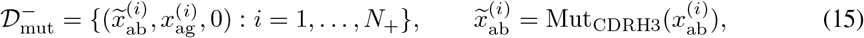

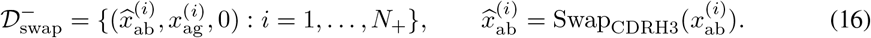

Here, 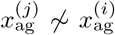 indicates that antigen 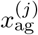 is sufficiently dissimilar to the cognate antigen 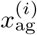. Mut_CDRH3_ generates a perturbed antibody by modifying the CDRH3 sequence, a region important for antibody–antigen binding [Forthal, 2014]. Swap_CDRH3_ replaces the original CDRH3 with a length-matched naturally occurring sequence from a reference CDRH3 library. These pseudo-negative sets represent distinct sources of specificity perturbation: antigen mismatch, local CDRH3 mutation, and CDRH3 replacement. During training, positive and pseudo-negative examples were sampled at a ratio of 1:2. Appendix E.4 reports the effects of different pseudo-negative types and sampling ratios on specificity prediction. Full construction details, including antigen similarity thresholds, mutation constraints, CDRH3 library construction, and sampling ratios, are provided in Appendix A.1.

#### Training objectives of FoldAbS-Px

Let *D*_spec_, *D*_aff_, and 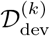 denote the training sets for specificity prediction, affinity prediction, and the *k*-th developability property, respectively. FoldAbS-Px was trained with task-specific objectives that matched the corresponding label formats. Since the Protenix encoder was kept fixed, only the downstream prediction-head parameters were optimized.

Specificity prediction was optimized with binary cross-entropy:

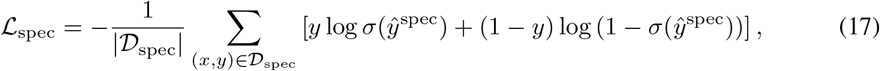

where *σ*(*·*) denotes the sigmoid function.

Affinity prediction was optimized using a regression loss over affinity labels:

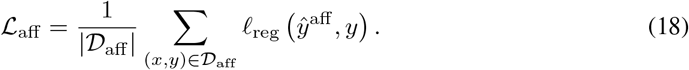

For developability prediction, each property was optimized separately:

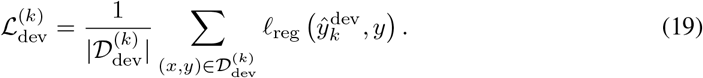

Here, *l*_reg_ denotes the regression loss used for affinity and developability prediction. Head architectures and optimization hyperparameters are provided in Appendix A.3.

## 4 Experiments

### 4.1 Specificity Prediction

#### Datasets

For specificity prediction, the training set was constructed from ASD, which contains 1,227,083 antibody–antigen pairs. After retaining entries with *high* or *very high* confidence, removing invalid sequence pairs, and restricting antigen lengths to 10–600 residues, the final training set contained 100,973 positive binding pairs. Since experimentally validated non-binding pairs are limited in ASD, pseudo-negative pairs were generated from these positives as described in Appendix A.1. During training, positive and pseudo-negative pairs were sampled at a 1:2 ratio.

Specificity prediction was evaluated on two main test sets. Origin-1 [Levine et al., 2026] integrated experimental antibody–antigen interaction data from two previous studies [Shanehsazzadeh et al., 2023a,b], comprising 3,099 pairs across eight antigen systems with 1,017 binders. Because most labels were derived from CDR mutagenesis, Origin-1 mainly tested specificity prediction under sequence variation around known antibody scaffolds. PPIFlow [Yu et al., 2026] contained experimental measurements for 30 de novo designed antibodies against each of seven targets, resulting in 210 pairs with 81 binders, and evaluated screening performance in a de novo design setting. These designed antibodies are substantially different from known database binders, making this benchmark a challenging test of generalization beyond known antibody scaffolds. A more challenging pair-shuffled cross-antigen benchmark [Smorodina et al., 2026] was additionally evaluated in Appendix E.2.

To reduce data leakage, the ASD training set was sequence-deduplicated separately for each test set. Antibody sequences from all positive test pairs were used as MMseqs2 queries, and ASD samples with sequence identity above 95% were removed [Steinegger and Söding, 2017]. The resulting train/test splits were therefore deduplicated with respect to positive pairs in each evaluation setting. Full data-processing details are provided in Appendix B.1.

#### Baselines and evaluation metrics

The specificity model was compared with both structure-based and language-model-based baselines. The structure-based baselines included Protenix-v1 [Zhang et al., 2026], the folding model used as the encoder of FoldAbS-Px, and AlphaFold 3 [Abramson et al., 2024]. For these two folding-model baselines, antibody–antigen pairs were ranked by the native interface confidence score, *ipTM*. The language-model-based baselines included ESM-650M [Lin et al., 2023], the antibody foundation model Ophiuchus-Ab [Zhu et al., 2026], and S3AI [Liu et al., 2024], which incorporates antibody sequence and structural information during pretraining. For a fair comparison, all learned baselines were fine-tuned on the ASD training set. Detailed implementation settings are provided in Appendix C.1.

For Origin-1, evaluation followed the protocol of prior work [Levine et al., 2026]. AUROC and enrichment factors at 1%, 5%, and 10% screening rates were reported, denoted as EF@1%, EF@5%, and EF@10%, respectively. For PPIFlow, AUROC was used as the primary metric because each target contains only 30 antibody candidates, making low-rate enrichment estimates sensitive to small changes in ranking. Enrichment factors were reported in Appendix E.3 for completeness and were consistent with the AUROC-based comparison.

#### Specificity prediction across benchmarks

Table 1 summarizes the specificity prediction results on the Origin-1 test set. In this benchmark, Protenix-v1 and AlphaFold 3 are evaluated as foldingmodel baselines using their native interface confidence score, ipTM. This setting provides a direct comparison between native folding-model confidence scores and the task-specific specificity head learned by FoldAbS-Px. FoldAbS-Px achieved the highest average AUROC of 0.757, exceeding Protenix-v1 and AlphaFold 3 by 0.088 and 0.104, respectively. At the antigen level, FoldAbS-Px outperformed Protenix-v1 on six of the eight antigen systems and also exceeded AlphaFold 3 on six of the eight systems, indicating that the improvement is not limited to a single target.

**Table 1:** Specificity performance on the Origin-1 test set. Best and second-best results in each column are shown in **bold** and underlined, respectively.

|  | AUROC |  |  |  |  |  |  |  |  | Early enrichment |  |  |
|---|---|---|---|---|---|---|---|---|---|---|---|---|
| Model | ACVR2B | C5 | FXI | HER2 | IL17A | IL36R | TNFRSF9 | TSLP | Avg. | EF@1% | EF@5% | EF@10% |
| *Folding-model-based methods* |  |  |  |  |  |  |  |  |  |  |  |  |
| FoldAbS-Px | 0.779 | **0.685** | **0.573** | 0.701 | **0.945** | **0.781** | 0.945 | 0.650 | **0.757** | **4.047** | **3.799** | **3.182** |
| Protenix-v1 | 0.610 | 0.681 | 0.479 | 0.380 | 0.922 | 0.620 | **0.965** | **0.691** | 0.669 | 3.872 | 2.052 | 2.374 |
| AF3 | **0.806** | 0.612 | 0.498 | 0.602 | 0.532 | 0.750 | 0.761 | 0.663 | 0.653 | 1.640 | 1.871 | 1.618 |
| *Language-model-based methods* |  |  |  |  |  |  |  |  |  |  |  |  |
| ESM-650M | 0.443 | 0.659 | 0.415 | 0.683 | 0.774 | 0.556 | 0.563 | 0.608 | 0.588 | 1.376 | 1.592 | 1.818 |
| Ophiuchus-Ab | 0.605 | 0.518 | 0.571 | **0.734** | 0.888 | 0.513 | 0.789 | 0.189 | 0.601 | 2.163 | 2.143 | 2.113 |
| S3AI | 0.443 | 0.481 | 0.534 | 0.614 | 0.592 | 0.379 | 0.241 | 0.394 | 0.460 | 1.081 | 0.747 | 0.649 |

Structure-based folding models consistently outperformed language-model-based baselines on this benchmark, highlighting the importance of explicitly modeling antibody–antigen geometry for specificity prediction. Beyond AUROC, FoldAbS-Px also achieved the best early enrichment across all screening rates. These results indicate that learning a task-aligned specificity head on top of Protenix improved over native folding-model confidence scores, leading to stronger binder discrimination and more effective prioritization of top-ranked candidates. Qualitative case studies comparing FoldAbS-Px scores with native Protenix ipTM are provided in Appendix E.6.

Table 2 reports specificity performance on the PPIFlow test set, which evaluates antibody screening in a de novo design setting. FoldAbS-Px achieved the best macro AUROC of 0.592, outperforming Protenix-v1 (0.559), AlphaFold 3 (0.525), and all language-model-based baselines. Relative to Protenix-v1, FoldAbS-Px achieved higher AUROC on five of seven targets, indicating that taskspecific supervision provided a more informative screening signal than native folding-model confidence scores. Relative to AlphaFold 3, FoldAbS-Px achieved higher AUROC on five of seven targets and remained competitive on the remaining two targets. These results suggest that FoldAbSPx transferred ASD-derived specificity supervision to experimentally validated de novo antibody candidates.

**Table 2:** Specificity AUROC on the PPIFlow test set. Best and second-best results are shown in **bold** and underlined, respectively.

| Model | 1433E | BHRF1 | CCL2 | EFNA1 | HNMT | PDGFR | S100A4 | Avg. |
|---|---|---|---|---|---|---|---|---|
| *Folding-model-based methods* |  |  |  |  |  |  |  |  |
| FoldAbS-Px | **0.833** | **0.816** | 0.448 | 0.545 | 0.326 | 0.656 | 0.520 | **0.592** |
| Protenix-v1 | 0.646 | 0.784 | 0.425 | 0.545 | 0.312 | **0.736** | 0.456 | 0.559 |
| AF3 | 0.549 | 0.792 | 0.371 | **0.582** | **0.639** | 0.384 | 0.360 | 0.525 |
| *Language-model-based methods* |  |  |  |  |  |  |  |  |
| ESM-650M | 0.444 | 0.616 | 0.181 | 0.376 | 0.354 | 0.384 | **0.536** | 0.413 |
| Ophiuchus-Ab | 0.535 | 0.632 | 0.344 | 0.344 | 0.271 | 0.520 | 0.432 | 0.440 |
| S3AI | 0.500 | 0.448 | **0.516** | 0.513 | 0.451 | 0.272 | 0.504 | 0.458 |

#### Comparison with interface-scoring metrics

We further compare FoldAbS-Px with ipSAE [Dunbrack Jr, 2025], pDockQ2 [Zhu et al., 2023], and actifpTM [Varga et al., 2025], computed from AF3 and Protenix-v1 outputs. As shown in Table 3, FoldAbS-Px achieves the highest macro AUROC on both benchmarks, exceeding the strongest interface-scoring baseline on Origin-1, AF3 actifpTM, by 0.0494 and the strongest baseline on PPIFlow, Protenix-v1 ipTM, by 0.0330. The relative performance of the interface scores varies across backbones and benchmarks: actifpTM improves AF3 on Origin-1, whereas none of the three additional metrics surpasses native ipTM for Protenix-v1 in macro AUROC. With the Protenix-v1 backbone held fixed, FoldAbS-Px increases macro AUROC from 0.6690 to 0.7570 on Origin-1 and from 0.5590 to 0.5920 on PPIFlow. These results demonstrate that supervised adaptation of folding-derived representations provides a stronger antibody-screening signal than the evaluated confidence and interface-scoring readouts. Additional language-model comparisons and evaluations with other folding backbones are reported in Appendices E.1 and E.7, respectively.

**Table 3:** Comparison with interface-scoring baselines for specificity prediction. We report macroaverage AUROC on Origin-1 and PPIFlow. Interface scores are computed from AF3 and Protenix-v1 outputs. Best and second-best results are shown in **bold** and underlined, respectively.

| Model | Scoring method | Origin-1 | PPIFlow |
|---|---|---|---|
| FoldAbS-Px | Learned specificity head | **0.7570** | **0.5920** |
| Protenix-v1 | ipTM | 0.6690 | 0.5590 |
|  | ipSAE | 0.6617 | 0.5348 |
|  | pDockQ2 | 0.6598 | 0.5442 |
|  | actifpTM | 0.6555 | 0.5162 |
| AF3 | ipTM | 0.6530 | 0.5250 |
|  | ipSAE | 0.6231 | 0.5211 |
|  | pDockQ2 | 0.6454 | 0.5068 |
|  | actifpTM | 0.7076 | 0.5135 |

### 4.2 Affinity Prediction

#### Datasets

For affinity prediction, FoldAbS-Px was evaluated on three benchmarks. The AFF and SKE benchmarks were adopted from Song et al. [2026], which provided standardized settings for antibody–antigen binding affinity prediction. AFF was constructed by combining affinity records from SAbDab [Dunbar et al., 2014] and BioMap [Chen et al., 2024], with 1,429 deduplicated antibody–antigen entries retained after preprocessing. The original AFF split contained 1,221/69/139 samples for training, validation, and testing, respectively. SKE was derived from SKEMPI 2.0 [Jankauskaitė et al., 2019] by retaining antibody–antigen mutation entries with quantitative affinity annotations. It contained 247 samples in total, with an original split of 214/11/22 samples for training, validation, and testing.

In addition to these public benchmarks, a HER2 affinity benchmark was constructed from the HER2 subset of Origin-1 by retaining antibody–antigen pairs with quantitative affinity annotations. The resulting 758 samples were split into 633/50/75 samples for affinity training, validation, and testing, respectively. This dataset provided an in-domain affinity evaluation setting closer to the antibody screening scenario studied in this work. Across all datasets, affinity labels were converted to the unified *pK*_*d*_ scale. Detailed dataset statistics, preprocessing procedures, and split information are provided in Appendix B.2.

#### Baselines and evaluation metrics

FoldAbS-Px was compared with general protein language models, antibody-specific language models, and task-specific affinity predictors. The language-model baselines included **ESM-650M** [Lin et al., 2023], **Ophiuchus-Ab** [Zhu et al., 2026], and **AntiBERTy** [Ruffolo et al., 2021]. The task-specific baselines included **EResCNN** [Gao et al., 2023], **A2binder** [He et al., 2024], and **AlphaBind** [Agarwal et al., 2024]. EResCNN was included as an ensemble residual convolutional baseline, while A2binder and AlphaBind were included as affinity-specific models pretrained on large-scale antibody–antigen affinity data. Additional comparisons across protein language model scales and families are reported in Appendix E.1. For a fair comparison, all learned baselines were fine-tuned on the corresponding training split of each dataset before evaluation. Detailed implementation settings are provided in Appendix C.2.

Following Song et al. [2026], **Spearman’s rank correlation coefficient** (*ρ*) and **mean absolute error** (MAE) were used as the primary evaluation metrics. Spearman’s *ρ* measures rank-level agreement between predicted and observed affinities, while MAE quantifies absolute prediction error in the unified *pK*_*d*_ space.

#### Affinity prediction across benchmarks

Table 4 summarizes affinity prediction performance on AFF-Test, SKE-Test, and HER2-Test. FoldAbS-Px achieved the best performance on all three benchmarks according to both Spearman’s *ρ* and MAE, with *ρ*/MAE values of 0.7406/0.9861, 0.9331/0.4882, and 0.7939/0.3055, respectively. The improvement was consistent across datasets with different characteristics: AFF-Test evaluated general antibody–antigen affinity prediction, SKE-Test focused on mutation-induced affinity changes, and HER2-Test provided an in-domain screeningoriented evaluation. Compared with the strongest competing baselines, FoldAbS-Px improved rank correlation and reduced absolute error. These results support the design choice of initializing the affinity head from a specificity-trained interaction module and fine-tuning it for continuous affinity regression. Joint training across standardized affinity datasets was further examined in Appendix E.5.

**Table 4:**
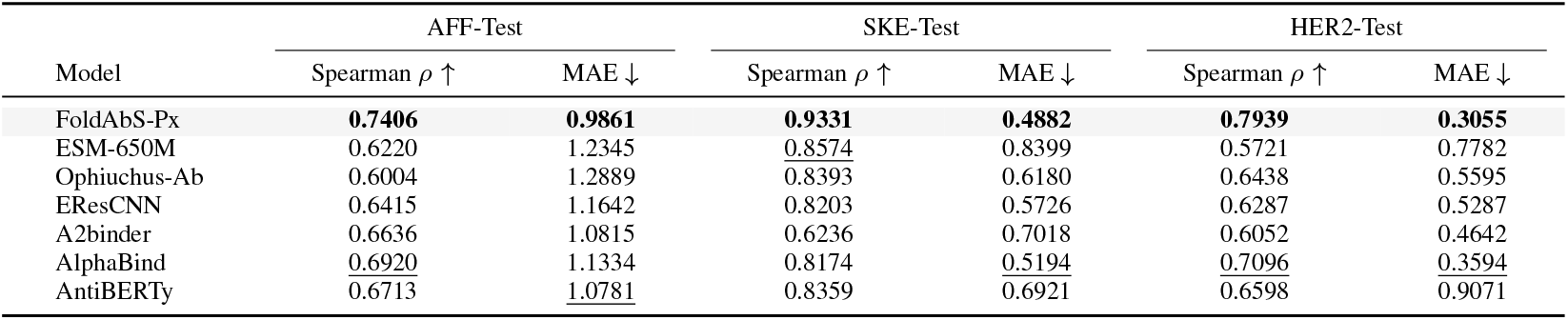
Affinity prediction performance across three test sets. Best and second-best results in each column are shown in **bold** and underlined, respectively.

|  | AFF-Test |  | SKE-Test |  | HER2-Test |  |
|---|---|---|---|---|---|---|
| Model | Spearman ρ ↑ | MAE ↓ | Spearman ρ ↑ | MAE ↓ | Spearman ρ ↑ | MAE ↓ |
| FoldAbS-Px | **0.7406** | **0.9861** | **0.9331** | **0.4882** | **0.7939** | **0.3055** |
| ESM-650M | 0.6220 | 1.2345 | 0.8574 | 0.8399 | 0.5721 | 0.7782 |
| Ophiuchus-Ab | 0.6004 | 1.2889 | 0.8393 | 0.6180 | 0.6438 | 0.5595 |
| EResCNN | 0.6415 | 1.1642 | 0.8203 | 0.5726 | 0.6287 | 0.5287 |
| A2binder | 0.6636 | 1.0815 | 0.6236 | 0.7018 | 0.6052 | 0.4642 |
| AlphaBind | 0.6920 | 1.1334 | 0.8174 | 0.5194 | 0.7096 | 0.3594 |
| AntiBERTy | 0.6713 | 1.0781 | 0.8359 | 0.6921 | 0.6598 | 0.9071 |

### 4.3 Developability Prediction

#### Datasets

For developability prediction, benchmarks were constructed based on FLAb2 [Chungyoun and Gray, 2025], a large-scale antibody developability resource that curates therapeutic antibody assay data across multiple biophysical properties. This resource was further augmented with a subset of in-house developability measurements. Four properties were considered in this study: aggregation, expression, immunogenicity, and thermostability.

Each property was treated as an independent regression task. After preprocessing and harmonization, the aggregation dataset contained 192/22/35 samples for training, validation, and testing, respectively; the expression dataset contained 622/34/117 samples; the immunogenicity dataset contained 160/20/32 samples; and the thermostability dataset contained 337/30/63 samples. Detailed preprocessing procedures are provided in Appendix B.3.

#### Baselines and evaluation metrics

FoldAbS-Px was compared with baselines spanning general protein language models, antibody-specific language models, and structure-aware antibody representations. The baselines included **ESM-650M** [Lin et al., 2023] as a general protein language model, **Ophiuchus-Ab** [Zhu et al., 2026] and **AntiBERTy** [Ruffolo et al., 2021] as antibody-specialized language models, and three IgFold-based variants [Ruffolo et al., 2023]: *IgFold bert, IgFold struc*, and *IgFold gt*. Additional comparisons across protein language model scales and families are reported in Appendix E.1. For a fair comparison, all baselines were fine-tuned on the training split of each developability property and evaluated on the corresponding test set. Detailed implementation settings are provided in Appendix C.3.

Following the FLAb2 evaluation protocol, **Spearman’s rank correlation coefficient** (*ρ*) was used as the primary metric to assess agreement between predicted and experimentally observed rankings across developability assays with different numerical scales.

#### Developability prediction across properties

Table 5 summarizes developability prediction performance across four antibody properties. FoldAbS-Px achieved the best Spearman’s *ρ* on all four tasks, reaching 0.406 on aggregation, 0.348 on expression, 0.367 on immunogenicity, and 0.845 on thermostability. Compared with the strongest competing baselines, FoldAbS-Px improved over Ophiuchus-Ab from 0.363 to 0.406 on aggregation, over IgFold bert from 0.328 to 0.348 on expression, over AntiBERTy from 0.329 to 0.367 on immunogenicity, and over Ophiuchus-Ab from 0.832 to 0.845 on thermostability. These results suggest that Protenix-derived structural representations transferred beyond interaction prediction to antibody-centered developability properties. Five-fold cross-validation results for all developability baselines are reported in Appendix E.8.

**Table 5:** Developability prediction performance across four antibody properties. Best and second-best results in each column are shown in **bold** and underlined, respectively.

| Model | Aggregation | Expression | Immunogenicity | Thermostability |
|---|---|---|---|---|
| FoldAbS-Px | **0.406** | **0.348** | **0.367** | **0.845** |
| ESM-650M | 0.362 | 0.311 | 0.202 | 0.784 |
| IgFold bert | 0.240 | <u>0.328</u> | 0.297 | 0.712 |
| IgFold struc | 0.147 | 0.126 | 0.179 | 0.484 |
| IgFold gt | 0.246 | 0.072 | 0.058 | 0.471 |
| AntiBERTy | 0.264 | 0.226 | <u>0.329</u> | 0.702 |
| Ophiuchus-Ab | <u>0.363</u> | 0.327 | 0.299 | <u>0.832</u> |

## 5 Conclusions and Limitations

We presented **FoldAbS**, a unified framework that repurposes protein folding models as frozen foundation encoders for supervised antibody screening, instantiated as **FoldAbS-Px** with Protenix. FoldAbS-Px achieved state-of-the-art performance across specificity, affinity, and developability benchmarks, outperforming folding-model, language-model, and task-specific baselines. FoldAbS is modular and does not require redesigning the folding backbone, allowing it to be applied as an add-on screening module for modern folding models. Current limitations include a focus on prediction and screening rather than closed-loop antibody design, as well as a partial reliance on curated pseudonegatives for specificity supervision. Future work should integrate the learned screening signals into hallucination-based or generative design pipelines and develop more realistic negative-sample construction strategies to reduce false-negative noise and improve screening robustness.

## A Additional Method Details

### A.1 Pseudo-negative Construction for Specificity Supervision

As described in the main text, specificity prediction is formulated as binary classification over binder and non-binder antibody–antigen pairs. However, the available paired interaction data are dominated by observed binders, whereas experimentally validated non-binders are scarce. To enable supervised training, we construct a set of pseudo-negative pairs from the observed positive set using three pseudo-negative construction strategies.

We denote the positive set by

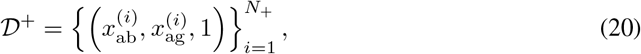

where 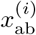 and 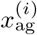 denote an antibody and its cognate antigen, respectively. The full pseudo-negative set is defined as

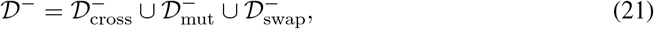

and the specificity training set is

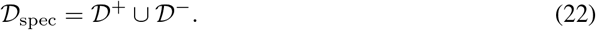

#### Cross-antigen negatives

The first pseudo-negative component is constructed by pairing a binder antibody with a non-cognate antigen whose sequence is sufficiently dissimilar to the cognate one. Formally,

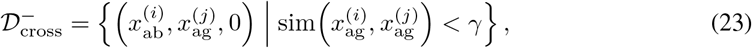

where sim(*·, ·*) denotes antigen sequence similarity and *γ* is a predefined threshold (set to 0.9 in our experiments). This strategy preserves the antibody while replacing its cognate antigen with a sufficiently different target, thereby creating coarse specificity mismatches.

#### Mutation-based negatives

The second pseudo-negative component perturbs the CDRH3 region of binder antibodies. Let

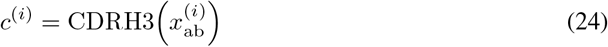

denote the CDRH3 sequence of antibody 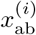. We generate a perturbed sequence 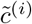 satisfying

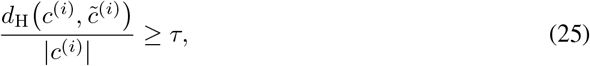

where *d*_H_(*·, ·*) is the Hamming distance, |*c*^(*i*)^ | denotes the CDRH3 length, and *τ* = 0.5 in our experiments. We further require

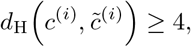

so that each perturbed CDRH3 contains at least four residue substitutions. Replacing the original CDRH3 by 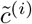 yields a perturbed antibody 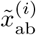, from which we form

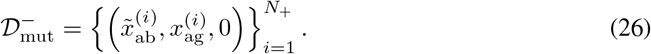

This strategy is motivated by the well-established role of CDRH3 as a dominant determinant of antibody–antigen recognition [Forthal, 2014]. Compared with cross-antigen negatives, mutationbased negatives preserve the overall antibody context while specifically perturbing a key recognition region.

#### Swap-based negatives

The third pseudo-negative component replaces the original CDRH3 with a naturally occurring sequence sampled from a reference CDRH3 library. For each binder sequence *c*^(*i*)^, we select a library sequence *c*^*′*^ *∈ C* satisfying

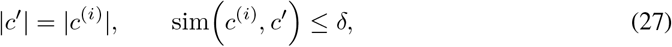

where *C* denotes the natural CDRH3 library and *δ* is a sequence-similarity threshold. Replacing *c*^(*i*)^ with *c*^*′*^ yields a swapped antibody 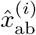, from which we define:

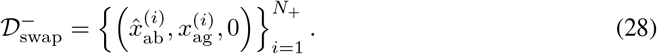

Compared with mutation-based negatives, this strategy produces harder decoys by preserving CDRH3 length and drawing replacement sequences from naturally occurring antibodies, while still disrupting antigen-specific recognition patterns.

#### Sampling and practical use

The three negative-generation strategies provide complementary supervision. Cross-antigen negatives represent coarse mismatched pairs, whereas mutation- and swapbased negatives produce harder decoys that remain closer to natural binders in antibody sequence space. In practice, these pseudo-negative pools are combined to form the training set at a positiveto-negative ratio of 1:2, as described in the main text. Within *D*^*−*^, the relative sampling ratio of cross-antigen, swap-based, and mutation-based negatives is set to 3:1:6. Additional implementation details, including the exact construction of the CDRH3 library and training-time cluster-based sampling of antibody sequences, are provided in our released code and scripts.

### A.2 Detailed Task-Head Architectures

This section provides implementation details for the task-specific heads used in FoldAbS-Px. In the main text, each task is written as

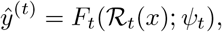

where ℛ_*t*_(*x*) denotes the Protenix-derived feature subset used by task *t*, and *ψ*_*t*_ denotes the trainable downstream parameters. Throughout all tasks, the Protenix encoder *P*_*θ*_ is frozen; only task-specific downstream modules are optimized.

Let 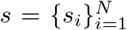 and 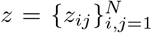 denote the single and pair representations produced by Protenix, and let 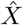, 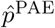, and 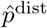 denote the predicted coordinates, confidence-related outputs, and distogram predictions, respectively. A common operation in the specificity and developability branches is pairwise conditioning, which injects task-dependent pairwise features into the Protenix pair representation:

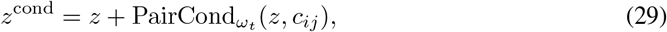

where *c*_*ij*_ is the task-specific pairwise condition feature. 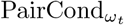 denotes a lightweight conditioning module implemented with layer normalization, linear projection, and residual transition blocks. Its parameters *ω*_*t*_ are a subset of the downstream head parameters *ψ*_*t*_, rather than the full task head. Algorithm S1 summarizes how the three task-specific heads consume Protenix-derived representations.

#### Specificity head

For specificity prediction, FoldAbS-Px uses the interaction-level feature set

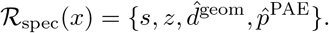

The geometry-derived distance feature 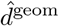 is obtained by discretizing pairwise distances between representative atoms in the predicted structure 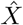 into 64 bins with maximum distance 22.0Å. The PAE logits are converted into probabilities and projected into the pair-representation space. These two pairwise signals are concatenated:

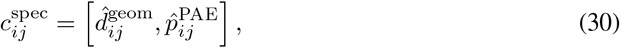

and injected into the pair representation through the pairwise conditioning module:

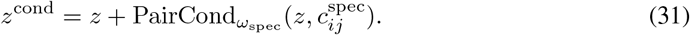

The conditioned pair representation is refined together with *s* using a 4-block task-specific Pairformer stack. We then pool cross-chain pair features and antibody/antigen single features to form an interfaceaware structural summary. The scalar iPTM score is embedded into a 32-dimensional vector and fused with this structural summary before the final classification MLP. The output is a binding logit *ŷ*^spec^. The pairwise conditioning parameters *ω*_spec_, Pairformer refinement layers, pooling layers, iPTM embedding, and classification MLP are all part of *ψ*_spec_.

##### Algorithm S1

Task-specific heads on top of frozen Protenix representations

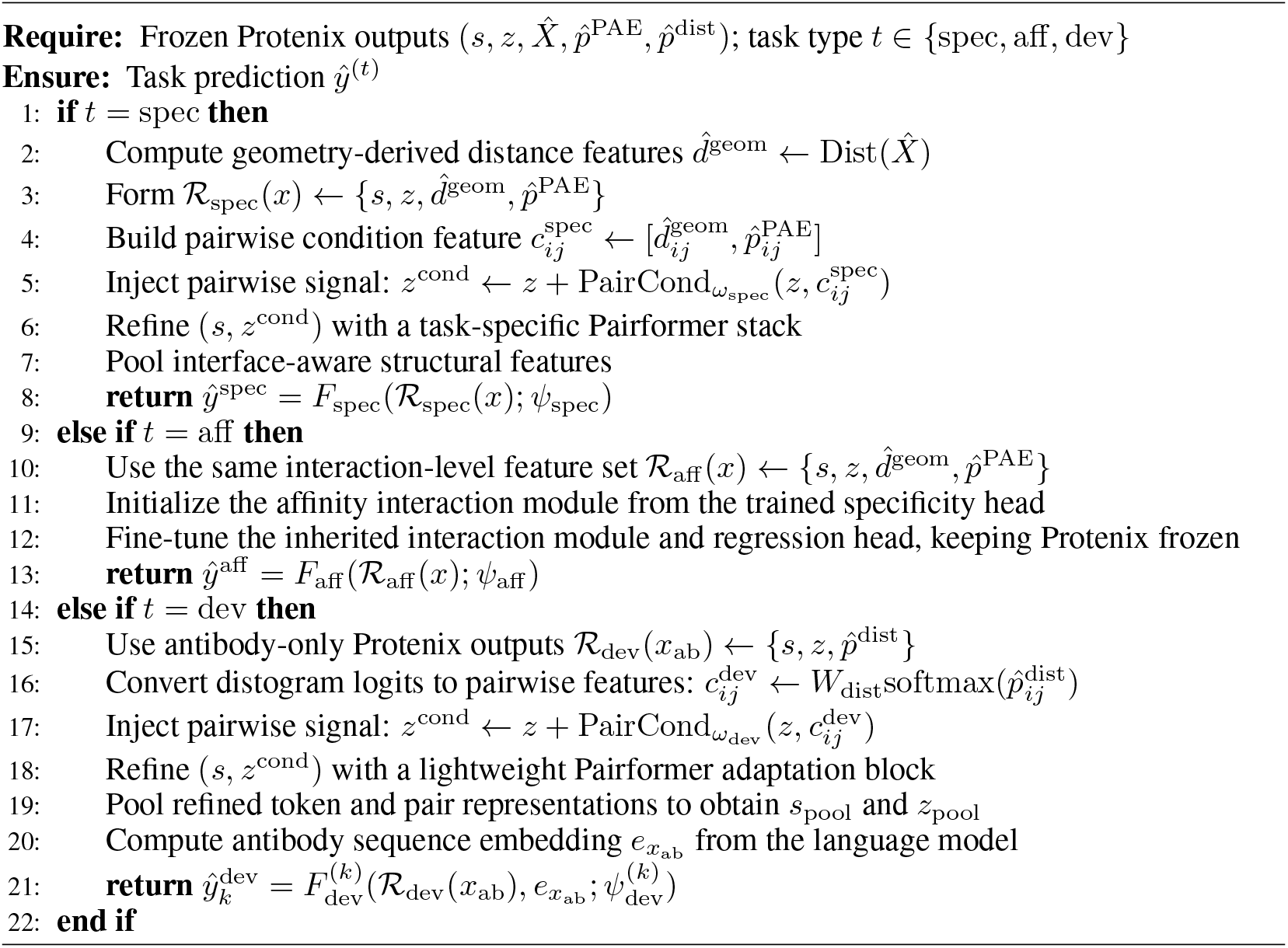

#### Affinity head

Affinity prediction uses the same interaction-level feature set as specificity:

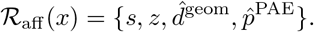

As in the main text, the affinity head is initialized from the specificity head trained for binder discrimination. Let 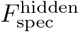 denote the specificity head up to its penultimate hidden layer, i.e., before the final binary classification layer. The affinity head replaces this final classification layer with a regression MLP:

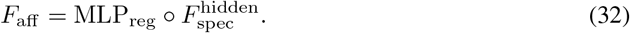

Thus, the affinity prediction is

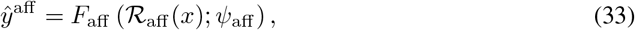

where *ψ*_aff_ includes the trainable parameters of both the inherited specificity module and the regression MLP.

During affinity training, the Protenix backbone remains frozen, while 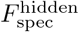 and MLP_reg_ are fine-tuned on quantitative binding labels. This design is important because affinity datasets are composed primarily of binding pairs: a frozen specificity head tends to map these examples to similarly high-confidence binder representations, whereas affinity prediction requires resolving finer differences in binding strength among binders. Fine-tuning the inherited specificity module adapts the representation from binary binder discrimination to continuous affinity regression. In the main configuration, MLP_reg_ is a 3-layer MLP with hidden dimension 128 and dropout 0.1.

#### Developability head

For developability prediction, Protenix is applied to the antibody input *x*_ab_. Since developability reflects antibody-intrinsic biophysical behavior more than antigen-specific interface geometry, FoldAbS-Px uses antibody-level features

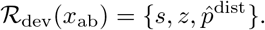

The distogram logits are converted into probabilities and projected into the pair-representation space:

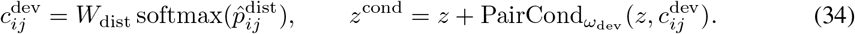

The conditioned pair representation and the single representation are then refined with a 1-block Pairformer adaptation module. After refinement, we compute pooled summaries

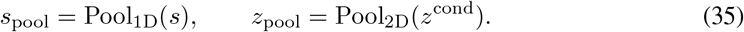

These structural summaries are concatenated with antibody sequence embeddings from frozen ESM-2:

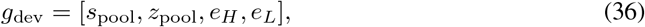

where *e*_*H*_ and *e*_*L*_ are mean-pooled heavy- and light-chain embeddings. A 3-layer MLP with hidden dimension 512 and dropout 0.1 predicts a scalar value for each developability property. Aggregation, expression, immunogenicity, and thermostability are modeled as separate single-property regression tasks with identical architecture but independent parameters. The pairwise conditioning parameters *ω*_dev_, Pairformer adaptation block, pooling operations, ESM-fusion projection, and property-specific MLP are included in 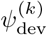.

#### Parameterization and implementation choices

Across all tasks, the Protenix backbone is initialized from the publicly available Protenix-v1 checkpoint, since the weights of Protenix-v2 are not released at the time of this study. The single and pair representation dimensions follow the AF3-style parameterization used by Protenix, with *c*_*s*_ = 384 and *c*_*z*_ = 128. For specificity, the pairwise conditioning module uses 64 geometry-distogram bins, 64 PAE bins, a maximum geometric distance 22.0Å, two transition layers, 32 diffusion samples in the mini-rollout, and 20 denoising steps. For developability, the pairwise conditioning module uses 64 distogram bins and two transition layers, followed by a 1-block Pairformer adaptation module. Frozen ESM-2 (esm2_t33_650M_UR50D) is used as the language-model feature extractor for the main developability model. Additional ablation variants are reported in the subsequent appendix sections.

### A.3 Training Objectives and Optimization Details

All downstream tasks followed the shared Protenix-based formulation described in the main text, but were trained separately with task-specific supervision and validation criteria. Across all experiments, the backbone was initialized from the publicly available Protenix-v1 checkpoint, because the weights of Protenix-v2 were not publicly available at the time of this study. Model selection was performed independently for each task using the corresponding validation set.

#### Specificity training

Specificity prediction was optimized as binary classification over binder and non-binder antibody–antigen pairs using binary cross-entropy with logits. Positive and pseudonegative pairs were sampled at a ratio of 1:2, and antibody sequences were further sampled with a cluster-aware strategy to reduce redundancy among highly similar binders. All trainable parameters were optimized using AdamW [Loshchilov and Hutter, 2017]. For the specificity head, the learning rate was set to 10^*−*4^, the batch size was set to 1, and gradients were accumulated over 3 steps. The model was trained for 4,000 optimization steps on 8 NVIDIA H20 GPUs. The trainable specificity head used 64 distogram bins, 64 PAE bins, a maximum geometric distance of 22.0Å, 32 diffusion samples in the mini-rollout, 20 denoising steps, and a 4-block Pairformer refinement stack.

#### Affinity training

Affinity prediction was optimized as scalar regression on top of the pooled interaction embedding inherited from the specificity head. In the main configuration reported in the main text, the inherited specificity interaction module was fine-tuned together with a lightweight regression head using mean squared error on affinity labels represented in the unified *pK*_*d*_ space. Trainable parameters were optimized using AdamW with a learning rate of 10^*−*4^, weight decay of 0.01, gradient accumulation over 3 steps, and gradient clipping with a maximum norm of 1.0. The model was trained for 4,000 optimization steps on 8 NVIDIA H20 GPUs. The affinity regression head was selected using validation MSE.

#### Developability training

Each developability property was trained as an independent scalar regression task. In the main configuration, both the Protenix backbone and the ESM-2 sequence encoder were frozen, and only the task-specific fusion head was optimized. Mean squared error was used as the training objective. Each developability model was optimized with AdamW using a learning rate of 10^*−*4^ and a weight decay of 0.01. Training was run for up to 5,000 steps on 8 NVIDIA H20 GPUs, with gradient accumulation over 3 steps, gradient clipping at 1.0, and validation every 100 steps. The learning rate was scheduled with 100 warmup steps, patience of 50, decay factor of 0.5, and a minimum learning rate of 10^*−*5^. The main developability head used a 1-block Pairformer refinement module, 64 distogram bins, two pairwise-conditioning transition layers, and a 3-layer MLP with hidden dimension 512 and dropout 0.1. Model checkpoints were selected independently for each developability property using the corresponding validation MSE.

## B Dataset Construction and Preprocessing

### B.1 Specificity Datasets

The specificity training data were derived from the Antigen-Specific Antibody Database (ASD) resource through a preprocessing pipeline designed to retain high-confidence antibody–antigen binding pairs and to prepare sequence annotations for negative construction and Protenix featurization.

#### ASD preprocessing and filtering

Heavy-chain, light-chain, and antigen sequences were normalized by uppercasing residues and removing whitespace. Sequences containing non-standard amino acids were discarded. Chain-consistency constraints were then applied: each entry was required to contain a valid heavy chain; non-nanobody antibodies were required to contain a light chain; and nanobody entries were required to lack a light chain. Only entries annotated as *high* or *very_high* confidence were retained. Boolean non-binding records were removed, and duplicate antibody– antigen triplets were collapsed by exact matching of heavy-chain, light-chain, and antigen sequences. Antibody–antigen pairs used in the test sets were excluded, and antigens were restricted to lengths between 10 and 600 residues.

#### Chothia annotation and sequence truncation

Antibody variable regions were annotated with Chothia numbering using ANARCI [Dunbar and Deane, 2016] to support CDRH3-based negative construction. For each chain, the CDR3 boundaries, the largest valid Chothia position, and the variable-region truncation index were recorded. Heavy chains were truncated at Chothia position 113 and light chains at position 106. These annotations defined the sequence regions used for mutation-based and swap-based CDRH3 perturbations.

#### Sequence clustering and deduplication

Antibody and antigen sequences were clustered with MMseqs2 [Steinegger and Söding, 2017]. Concatenated heavy/light antibody sequences were clustered at 95% sequence identity to define <monospace>antibody_cluster_id</monospace>, whereas antigen sequences were clustered at 90% identity to define <monospace>antigen_cluster_id</monospace>. For leakage reduction on external benchmarks, ASD training antibodies were jointly clustered with test antibodies, and ASD entries sharing a test-antibody cluster were removed.

After preprocessing, the final ASD training set contained 100,973 antibody–antigen pairs, grouped into 8,520 antibody clusters and 1,063 antigen clusters (Table S1). The final ASD training set remained long-tailed: a few antigens, including hemagglutinin and VEGF, dominated the antibodycluster distribution, whereas most antigens had limited antibody diversity. The 10 largest antigen clusters by distinct antibody-cluster count are reported in Table S2, motivating cluster-balanced sampling and cross-antigen negative construction.

**Table S1:** Cluster counts for the final ASD training set after preprocessing.

| Quantity | Count |
|---|---|
| Total antibody–antigen pairs | 100,973 |
| Unique antibody clusters (antibody_cluster_id) | 8,520 |
| Unique antigen clusters (antigen_cluster_id) | 1,063 |

#### Training-time sampling

Specificity training used an iterable dataset organized by antibody cluster rather than by raw row. At each epoch, antibody clusters were shuffled, and one positive binder pair was randomly sampled from each cluster. Two pseudo-negative examples were generated for each positive example, giving a 1:2 positive-to-negative ratio. The negative pool consisted of cross-antigen mismatches, CDRH3 mutation-based negatives, and natural CDRH3 swap negatives, as detailed in Appendix A.1. In the main experiments, mutation-based negatives used a CDRH3 mutation rate of 0.5 with at least four substitutions, and the dataloader controlled the mixture of antibody-swap and CDRH3-perturbation negatives. Validation examples were generated once through a deterministic cache so that the same pseudo-negative pairs were used across runs.

**Table S2:** Top-10 antigen clusters in the ASD training set ranked by the number of associated distinct antibody clusters. ag_clst denotes the antigen cluster identifier, and #Ab clusters / #Antibodies report the numbers of distinct antibody clusters and distinct heavy-light antibody pairs associated with each antigen, respectively.

| Rank | Target name | ag_clst | Seq. len. | #Ab clusters | #Antibodies |
|---|---|---|---|---|---|
| 1 | HA, hemagglutinin (Influenza) | 409 | 519 | 2,443 | 65,536 |
| 2 | VEGF, short isoform | 50 | 102 | 1,293 | 2,228 |
| 3 | VEGF, long isoform | 50 | 395 | 568 | 4,283 |
| 4 | – (AAYL49 nanobody library) | 374 | 14 | 538 | 17,585 |
| 5 | Rabies lyssavirus glycoprotein | 77 | 524 | 136 | 146 |
| 6 | CSP, circumsporozoite (Malaria) | 638 | 412 | 95 | 180 |
| 7 | Ricin | 649 | 576 | 53 | 57 |
| 8 | GFP, green fluorescent protein | 241 | 238 | 49 | 51 |
| 9 | HA, hemagglutinin (Influenza) | 655 | 518 | 48 | 1,917 |
| 10 | Interferon-γ | 70 | 166 | 45 | 48 |

#### Antigen cropping

To reduce memory cost and improve training efficiency, antigens longer than 256 residues were randomly cropped during training to a contiguous 256-residue window. The crop position was sampled uniformly for each example, and the corresponding precomputed antigen MSA was cropped to the same residue interval before featurization. Cropping was disabled during validation and testing, for which full antigen sequences were used.

#### MSA and structure-related feature construction

Precomputed MSAs were used for all antibody and antigen chains. To reduce preprocessing cost, antibody MSAs were searched only against UniRef50, whereas antigen MSAs followed the standard Protenix MSA search pipeline. The processed antibody and antigen sequences, together with their MSA directories, were then converted into Protenix input features through the standard inference pipeline. For mutation- and swap-based pseudo-negatives, the antibody MSA from the corresponding parent positive example was reused rather than recomputed for the perturbed sequence, while the antigen MSA remained unchanged. This choice improved efficiency and avoided introducing additional variation from MSA retrieval. Template usage was disabled in the specificity dataloader, with dummy template features inserted when required. The resulting feature dictionary contained token-level features, atom-to-token mappings, representative-atom masks, and affinity-token masks for the downstream Protenix-based specificity model.

### B.2 Affinity Datasets

Affinity datasets were processed with the same Protenix feature-construction pipeline used for specificity modeling, including precomputed MSAs and conversion to Protenix input features through the standard inference pipeline (Appendix B.1). This section, therefore, focuses on the construction and filtering of affinity supervision data.

#### AFF benchmark

Following InterAb [Song et al., 2026], the affinity benchmark was constructed by combining antibody–antigen affinity data from SAbDab [Dunbar et al., 2014] and BioMap [Chen et al., 2024]. The SAbDab subset contained experimentally characterized antibody–antigen complexes with quantitative binding measurements, whereas the BioMap subset was derived from the 2021 Global Antibody Affinity Prediction Challenge. After source merging and duplicate removal, the resulting AFF dataset contained 1,429 samples, with 1,221/69/139 samples assigned to training, validation, and test sets following the InterAb split. Sequence fields were further standardized, invalid entries were removed, and all affinity labels were converted to a unified logarithmic *pK*_*d*_ scale.

#### SKE benchmark

Following InterAb, the SKE benchmark was derived from SKEMPI 2.0 [Jankauskaitė et al., 2019] by extracting antibody–antigen mutation entries with quantitative affinity annotations. This dataset provided a complementary setting for evaluating mutation-associated affinity variation. After filtering invalid entries, the resulting SKE dataset contained 247 samples, with 214/11/22 samples assigned to training, validation, and test sets under the InterAb split. All labels were converted to the unified *pK*_*d*_ scale before training and evaluation.

#### HER2 benchmark

A HER2 affinity dataset was additionally constructed from the HER2 subset of the Origin-1 benchmark used in the specificity experiments. Only antibody–antigen pairs with quantitative affinity annotations were retained, yielding 758 samples split into 633/50/75 training, validation, and test samples. This dataset provided a screening-oriented affinity benchmark aligned with the main experimental setting. To reduce train/test leakage, HER2-related training entries were filtered using the same sequence-deduplication protocol described in Appendix B.1. All affinity labels were converted to the common *pK*_*d*_ scale.

#### Label standardization

The three affinity benchmarks provided different original label types and were mapped to a unified *pK*_*d*_ scale. For AFF, binding free energies Δ*G* in kcal/mol were converted as

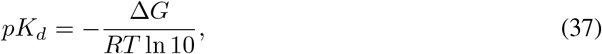

where *R* denotes the gas constant and *T* was set to 298.15 K. For SKE, labels were already provided as *pK*_*d*_ values, defined as *−* log_10_ *K*_*D*_ with *K*_*D*_ in molar units, and therefore required no transformation. For HER2, reported dissociation constants in nanomolar units were converted by

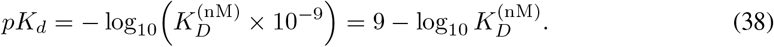

All affinity models were then trained and evaluated as regression models in the unified *pK*_*d*_ space. Dataset sizes and label statistics before and after conversion are summarized in Table S3.

### B.3 Developability Datasets

**Table S3:** Per-split sample counts and label statistics (mean and standard deviation) for the three affinity benchmarks, reported on both the original scale and the transformed *pK*_*d*_ scale. AFF is provided as Δ*G* (kcal/mol), HER2 as *K*_*D*_ (nM), and SKE is natively in *pK*_*d*_ (shown in the *pK*_*d*_ columns only).

| Dataset | Split | *N* | Original scale |  | *pK<sub>d</sub>* scale |  |
|---|---|---|---|---|---|---|
|  |  |  | Mean | Std | Mean | Std |
| AFF (Δ*G*, kcal/mol) | Train | 1,221 | −10.87 | 2.35 | 7.97 | 1.72 |
|  | Val | 69 | −10.59 | 2.13 | 7.76 | 1.56 |
|  | Test | 139 | −10.47 | 2.68 | 7.68 | 1.97 |
|  | All | 1,429 | −10.82 | 2.38 | 7.93 | 1.74 |
| SKE (native *pK<sub>d</sub>*) | Train | 214 | – | – | 7.81 | 1.27 |
|  | Val | 11 | – | – | 8.20 | 0.85 |
|  | Test | 22 | – | – | 7.95 | 1.07 |
|  | All | 247 | – | – | 7.84 | 1.24 |
| HER2 (*K<sub>D</sub>*, nM) | Train | 633 | 116.86 | 250.93 | 7.42 | 0.64 |
|  | Val | 50 | 163.81 | 355.75 | 7.45 | 0.77 |
|  | Test | 75 | 161.77 | 378.91 | 7.34 | 0.67 |
|  | All | 758 | 124.40 | 274.64 | 7.42 | 0.65 |

Four developability tasks were constructed for aggregation, expression, immunogenicity, and thermostability by combining assay-consistent FLAb2 subsets [Chungyoun and Gray, 2025] with selected in-house measurements. Each task was restricted to one assay family to reduce cross-study heterogeneity: SEC-based aggregation, HEK/HEK293 expression titer, clinical %ADA response, and DSF-derived melting temperature. In-house full-length antibodies were truncated to Fv regions using ANARCI [Dunbar and Deane, 2016] with Chothia numbering, whereas Fv-level FLAb sequences were used directly. Repeated heavy/light-chain pairs were merged by median assay value. Aggregation and immunogenicity labels were log(1 + *x*) transformed, and all task labels were standardized using train-split statistics. Splits were generated at the unique heavy-chain level. Dataset composition and label statistics are summarized in Tables S4 and S5.

**Table S4:**
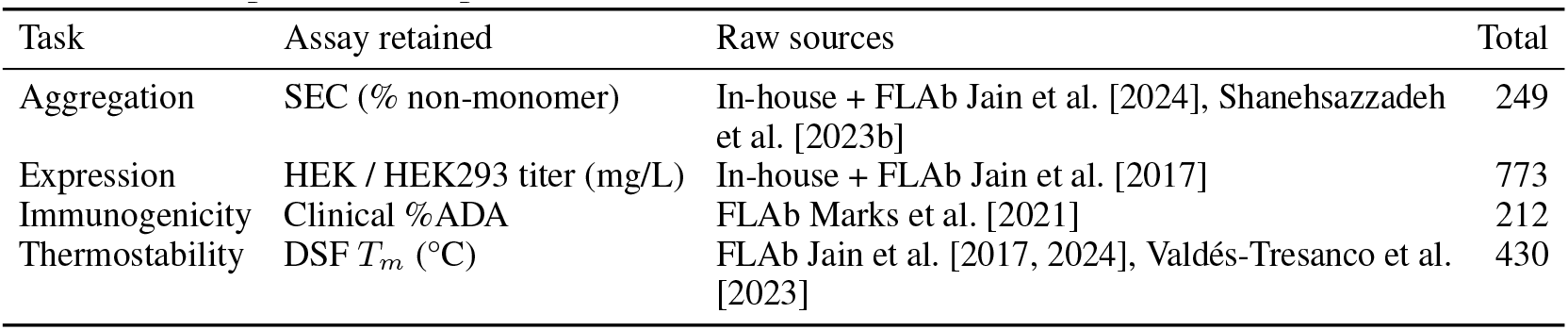
Developability benchmark composition before and after preprocessing. For each task, we report the retained assay family, raw source composition, the final number of samples after Fv truncation, and per-task deduplication.

**Table S5:** Training-split label statistics for the developability tasks. Raw labels are reported as mean *±* standard deviation in their native units. Processed labels denote the values used for model training after task-specific transformation and train-split z-score normalization. Aggregation and immunogenicity were first transformed by log(1 + *x*), whereas expression and thermostability were normalized directly from raw values.

| Task | Raw train labels | Processed train labels | Transform |
|---|---|---|---|
| Aggregation | 4.32 ± 7.89 (% non-monomer) | 0 ± 1 | log(1 + *x*) + z-score |
| Expression | 134.0 ± 57.5 (mg/L) | 0 ± 1 | z-score |
| Immunogenicity | 10.4 ± 18.2 (%ADA) | 0 ± 1 | log(1 + *x*) + z-score |
| Thermostability | 69.1 ± 9.0 (°C) | 0 ± 1 | z-score |

### C Experimental Setup and Baseline Details

### C.1 Specificity Baselines

Two groups of baselines were evaluated for specificity prediction: structure-based folding models and language-model-based predictors. Unless otherwise specified, all learned baselines were trained on the same ASD-derived training set as FoldAbS-Px and used the same positive-to-negative ratio.

#### Structure-based folding-model baselines

For **Protenix-v1** [Zhang et al., 2026] and **AlphaFold 3** [Abramson et al., 2024], the native folding-model confidence score *iPTM* was used as the screening score. Antibody and antigen MSAs were enabled during inference. For each antibody–antigen pair, diffusion inference was performed with 200 denoising steps and 25 structural hypotheses, generated from 5 fixed random seeds {42, 52, 62, 72, 82} and 5 samples per seed. The maximum *iPTM* among the 25 predictions was used as the final ranking score. This protocol assessed the screening utility of native folding-model confidence scores under multi-sample inference.

#### Language-model-based baselines

For **ESM-650M** [Lin et al., 2023], **Ophiuchus-Ab** [Zhu et al., 2026], and **S3AI** [Liu et al., 2024], antibody and antigen embeddings were extracted from the corresponding pretrained model, pooled into fixed-length representations, and passed to a lightweight specificity classification head. These predictors were trained on the ASD training set using the same specificity supervision protocol and positive-to-negative sampling ratio as FoldAbS-Px. Unless otherwise specified, all language-model baselines were trained for 5,000 optimization steps with a learning rate 10^*−*4^, weight decay 0.01, and batch size 32.

All learned specificity baselines were optimized with the same binary classification objective as FoldAbS-Px. A shared fine-tuning protocol was used to reduce confounding from baseline-specific hyperparameter tuning and to focus the comparison on the underlying pretrained representations.

#### Inference runtime

We measure wall-clock inference time on all 3,099 Origin-1 pairs using eight NVIDIA H20 GPUs, with the same MSA features, 10 trunk recycling cycles, 200 diffusion steps, and five samples per seed for both methods. Across five seeds executed sequentially, Protenix-v1 and FoldAbS-Px require 113.73 *±* 0.26 and 126.14 *±* 0.45 minutes per seed, respectively (mean *±* standard deviation). FoldAbS-Px thus improves screening performance with an additional 12.41 minutes per seed, corresponding to a 10.91% increase over the underlying folding inference.

### C.2 Affinity Baselines

For affinity prediction, FoldAbS-Px was compared with baselines covering general protein language models, antibody-specific language models, and task-specific affinity predictors, including **ESM650M** [Lin et al., 2023], **Ophiuchus-Ab** [Zhu et al., 2026], **AntiBERTy** [Ruffolo et al., 2021], **A2binder** [He et al., 2024], **EResCNN** [Gao et al., 2023], **and AlphaBind** [Agarwal et al., 2024].

ESM-650M, Ophiuchus-Ab, AntiBERTy, and A2binder were evaluated under a unified protocol. Antibody and antigen sequences were encoded with the corresponding pretrained model, pooled into fixed-length representations, concatenated, and passed to a lightweight regression head. Each model was fine-tuned separately on AFF, SKE, and HER2 using a learning rate of 10^*−*4^, batch size 32, and 5,000 training steps, with checkpoints selected by validation performance.

For EResCNN and AlphaBind, the official fine-tuning pipelines were used. Both models were retrained on the same affinity splits as FoldAbS-Px, and the checkpoint with the best validation performance was reported. This setup adapted all baselines to the same supervision setting while preserving architecture-specific training procedures for methods with released fine-tuning code.

### C.3 Developability Baselines

Following the FLAb2 benchmark [Chungyoun and Gray, 2025], FoldAbS-Px was compared with baselines covering general protein language models, antibody-specific language models, and structure-aware antibody representations. The baseline suite included **ESM-650M** [Lin et al., 2023], **Ophiuchus-Ab** [Zhu et al., 2026], **AntiBERTy** [Ruffolo et al., 2021], and three **IgFold**-based variants [Ruffolo et al., 2023]: *IgFold bert, IgFold struc*, and *IgFold gt*.

All baselines were evaluated under a unified developability prediction protocol. Antibody sequences, or the corresponding pretrained-model representations, were encoded and pooled into fixed-length embeddings, followed by a lightweight multilayer perceptron for property regression. For the IgFold variants, *IgFold bert* used the sequence-language branch, *IgFold struc* used the structure-derived representation, and *IgFold gt* used representations from ground-truth structures.

Each baseline was trained independently on the training split of each property and evaluated on the corresponding test split. Unless otherwise specified, all baseline fine-tuning runs used a batch size of 32, a learning rate of 10^*−*4^, and 3,000 training steps. This shared protocol provided a controlled comparison across developability properties and reduced confounding from baselinespecific hyperparameter tuning.

### C.4 Evaluation Metrics

Task-specific metrics were used according to the prediction target. Specificity was evaluated as a screening and ranking task, affinity as continuous regression, and developability as property-wise rank correlation.

#### Specificity metrics

For specificity prediction, AUROC was used as the primary ranking metric, measuring whether positive antibody–antigen pairs received higher scores than negative pairs. For screening-oriented evaluations, enrichment factors (EFs) were additionally reported at selected screening rates. Given a screening fraction *α*, the enrichment factor is defined as

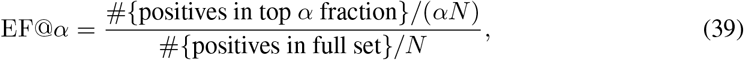

where *N* is the total number of candidate pairs. Intuitively, EF@*α* measures how much the model enriches true binders among the top-ranked fraction of candidates relative to random selection. In the main text, we report EF@1%, EF@5%, and EF@10% when the test-set size is sufficient for stable estimation.

#### Affinity metrics

For affinity prediction, we use Spearman’s rank correlation coefficient (*ρ*) and mean absolute error (MAE). Spearman’s *ρ* is defined as the Pearson correlation between the ranktransformed predictions and targets,

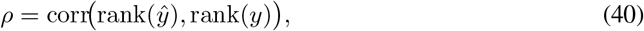

and measures whether the predicted ordering of affinity values agrees with the experimentally observed ordering. MAE is defined as

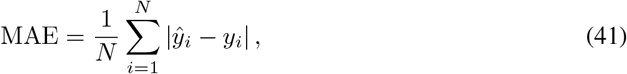

and quantifies absolute prediction error in the unified *pK*_*d*_ space.

**Table S6:**
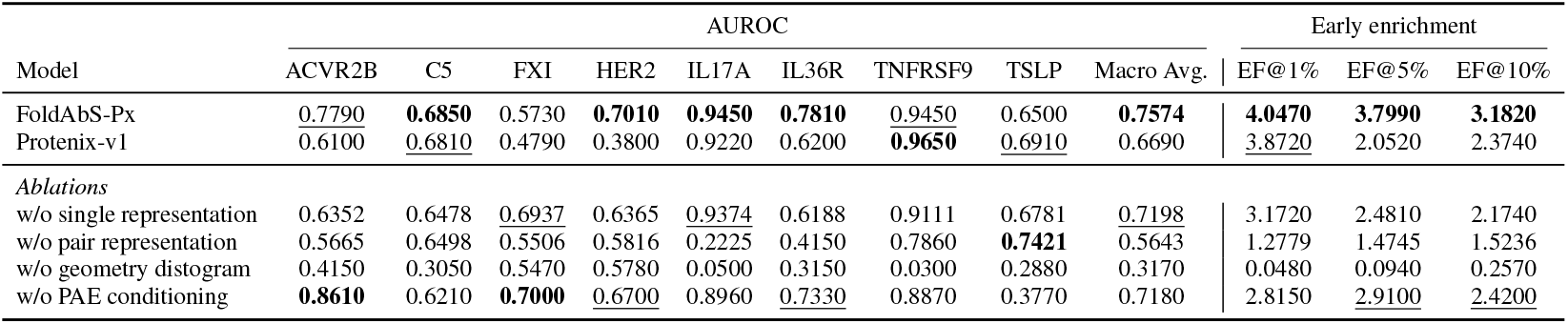
Specificity head ablations on Origin-1. We compare the full FoldAbS-Px model, native Protenix-v1 ipTM ranking, and four ablated specificity heads. Singleand pair-representation ablations remove direct access to the corresponding trunk features at the task-head input. Best and second-best results in each column are shown in **bold** and underlined, respectively.

|  | AUROC |  |  |  |  |  |  |  |  | Early enrichment |  |  |
|---|---|---|---|---|---|---|---|---|---|---|---|---|
| Model | ACVR2B | C5 | FXI | HER2 | IL17A | IL36R | TNFRSF9 | TSLP | Macro Avg. | EF@1% | EF@5% | EF@10% |
| FoldAbS-Px | 0.7790 | **0.6850** | 0.5730 | **0.7010** | **0.9450** | **0.7810** | 0.9450 | 0.6500 | **0.7574** | **4.0470** | **3.7990** | **3.1820** |
| Protenix-v1 | 0.6100 | 0.6810 | 0.4790 | 0.3800 | 0.9220 | 0.6200 | **0.9650** | 0.6910 | 0.6690 | 3.8720 | 2.0520 | 2.3740 |
| *Ablations* |  |  |  |  |  |  |  |  |  |  |  |  |
| w/o single representation | 0.6352 | 0.6478 | 0.6937 | 0.6365 | 0.9374 | 0.6188 | 0.9111 | 0.6781 | 0.7198 | 3.1720 | 2.4810 | 2.1740 |
| w/o pair representation | 0.5665 | 0.6498 | 0.5506 | 0.5816 | 0.2225 | 0.4150 | 0.7860 | **0.7421** | 0.5643 | 1.2779 | 1.4745 | 1.5236 |
| w/o geometry distogram | 0.4150 | 0.3050 | 0.5470 | 0.5780 | 0.0500 | 0.3150 | 0.0300 | 0.2880 | 0.3170 | 0.0480 | 0.0940 | 0.2570 |
| w/o PAE conditioning | **0.8610** | 0.6210 | **0.7000** | 0.6700 | 0.8960 | 0.7330 | 0.8870 | 0.3770 | 0.7180 | 2.8150 | 2.9100 | 2.4200 |

#### Developability metrics

For developability prediction, we report Spearman’s rank correlation coefficient (*ρ*) separately for each property. Since the developability tasks are formulated as singleproperty regression problems and the absolute scales differ substantially across properties, rank-based evaluation provides a stable assessment of whether a model correctly orders antibodies by the target biophysical attribute. Because Spearman’s *ρ* depends only on rank ordering, it is invariant to the monotonic preprocessing transforms used in our pipeline (e.g., log(1 + *x*) and z-score normalization); therefore, the same value is obtained whether the metric is computed in the transformed space or after mapping predictions back to the original property scale.

## D Ablations

### D.1 Specificity Ablations

We ablate the trunk representations and auxiliary structural signals used by the specificity head to assess their contributions to binder discrimination. The full model combines Protenix single and pair representations with a geometry-derived distogram, PAE features, and the scalar ipTM score. We compare four ablated variants: *w/o single representation, w/o pair representation, w/o geometry distogram*, and *w/o PAE conditioning*.

For the singleand pair-representation ablations, the corresponding tensor is zeroed only at the input to the task-specific head, and the ablated head is retrained. The original representations are retained inside frozen Protenix, preserving its structure-generation pathway. The geometry-distogram ablation removes the coordinate-derived distance features from pairwise conditioning. The PAE-conditioning ablation jointly removes the PAE features and scalar ipTM injection. All variants are evaluated on Origin-1 using per-antigen AUROC, macro-average AUROC, and early enrichment factors.

Table S6 identifies decoded geometry and direct pair-trunk access as the primary contributors to specificity performance. Removing the geometry-derived distogram reduces macro AUROC from 0.7574 to 0.3170 and sharply degrades early enrichment. Removing direct access to the pair representation also causes a substantial loss, reducing macro AUROC to 0.5643 and EF@10% from 3.1820 to 1.5236. These results establish the importance of combining explicit predicted geometry with the learned pairwise features carried by the folding-model trunk.

Single representations and confidence features provide complementary gains. Removing the single representation reduces macro AUROC to 0.7198, while removing PAE conditioning and ipTM injection reduces it to 0.7180. Both variants retain stronger macro AUROC than native Protenix-v1 ipTM ranking, but underperform the full model in all three enrichment metrics. The full model achieves the highest macro AUROC and early enrichment across all screening rates, demonstrating the benefit of integrating trunk representations, decoded geometry, and confidence information for antibody screening.

**Table S7:** Effect of inference sampling on Origin-1. Sampling configurations are denoted as the number of seeds *×* diffusion samples per seed. We report macro-average AUROC and enrichment factors. Best results within each model group are shown in **bold**.

| Model | Seeds × samples | Macro AUROC | EF@1% | EF@5% | EF@10% |
|---|---|---|---|---|---|
| FoldAbS-Px | 1 × 5 | 0.739 | 3.631 | 2.810 | 2.717 |
|  | 5 × 5 (default) | **0.757** | 4.047 | **3.799** | **3.182** |
|  | 10 × 5 | **0.757** | **4.172** | 3.551 | 3.018 |
| Protenix-v1 | 1 × 5 | 0.667 | 2.007 | 2.700 | 2.066 |
|  | 5 × 5 (default) | 0.669 | **3.872** | 2.052 | **2.374** |
|  | 10 × 5 | **0.673** | 3.267 | **2.936** | 2.197 |

#### Effect of inference sampling

We evaluate the effect of inference sampling by varying the number of random seeds while retaining five diffusion samples per seed. The default configuration uses five seeds, yielding 25 structural hypotheses per antibody–antigen pair. Table S7 compares one, five, and ten seeds on Origin-1 for FoldAbS-Px and native Protenix-v1 ipTM ranking.

Increasing the number of seeds from one to five improves FoldAbS-Px macro AUROC from 0.739 to 0.757 and increases all three enrichment factors. Further increasing to ten seeds leaves macro AUROC unchanged, slightly improves EF@1%, and reduces EF@5% and EF@10%. Native Protenix-v1 exhibits only a small increase in macro AUROC with additional seeds, while its enrichment factors vary non-monotonically. FoldAbS-Px outperforms native ipTM in macro AUROC at every matched sampling budget. These results support the default five-seed configuration, which achieves the same macro AUROC as ten seeds with half the structural sampling budget and stronger enrichment at the 5% and 10% screening rates.

### D.2 Affinity Ablations

The role of specificity pretraining in downstream affinity regression was examined by ablating how the affinity head used the interaction representation inherited from the specificity head. In all variants, the Protenix backbone was kept frozen.

As described in Appendix A.2, the affinity head is written as

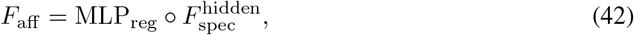

where 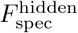 denotes the specificity head up to its penultimate hidden layer and MLP_reg_ is the final affinity regression head. The affinity prediction is therefore

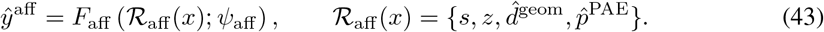

Four variants were compared. In the *frozen specificity head* setting, 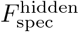 was initialized from the specificity-trained model and kept fixed, and only the regression MLP MLP_reg_ was trained. This setting tested whether the specificity-derived hidden representation was sufficient for affinity regression without task-specific adaptation.

In the *unfrozen specificity head* setting, 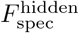 was initialized from the specificity-trained model but fine-tuned together with MLP_reg_. This is the main affinity model used in the paper. Since affinity datasets consist primarily of binding antibody–antigen pairs, a frozen specificity module tends to map examples to similarly high-confidence binder embeddings. Fine-tuning 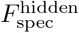 adapts the representation from binary binder discrimination to continuous differences in binding strength.

In the *+ ESM sequence fusion* setting, the unfrozen affinity model was augmented by concatenating frozen ESM-derived antibody and antigen embeddings with the specificity-derived hidden representation:

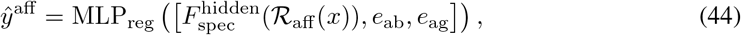

where *e*_ab_ and *e*_ag_ denote pooled antibody and antigen sequence embeddings, respectively. This variant tests whether additional language-model sequence information complements the Protenixderived interaction representation.

**Table S8:**
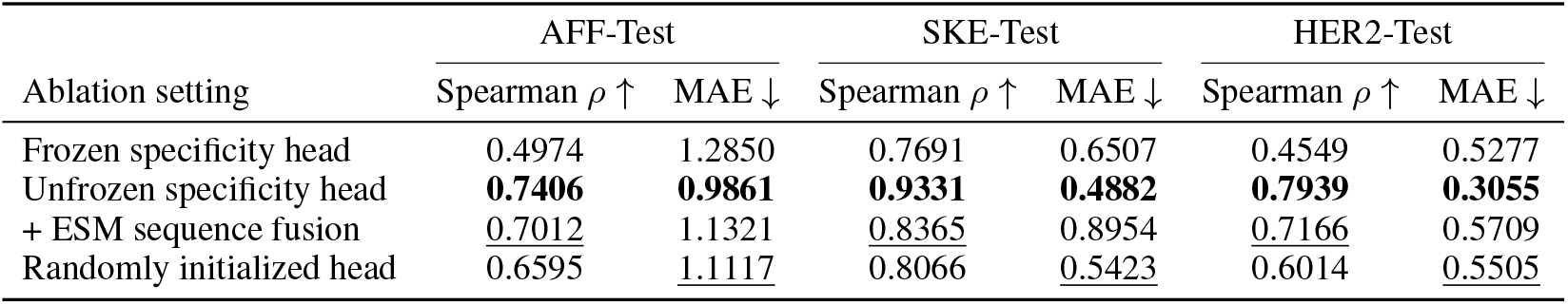
Ablation study of affinity prediction. We report Spearman’s rank correlation coefficient (*ρ*) and mean absolute error (MAE) on AFF-Test, SKE-Test, and HER2-Test. Best and second-best results in each column are shown in **bold** and underlined, respectively.

| Ablation setting | AFF-Test |  | SKE-Test |  | HER2-Test |  |
|---|---|---|---|---|---|---|
|  | Spearman ρ ↑ | MAE ↓ | Spearman ρ ↑ | MAE ↓ | Spearman ρ ↑ | MAE ↓ |
| Frozen specificity head | 0.4974 | 1.2850 | 0.7691 | 0.6507 | 0.4549 | 0.5277 |
| Unfrozen specificity head | **0.7406** | **0.9861** | **0.9331** | **0.4882** | **0.7939** | **0.3055** |
| + ESM sequence fusion | 0.7012 | 1.1321 | 0.8365 | 0.8954 | 0.7166 | 0.5709 |
| Randomly initialized head | 0.6595 | 1.1117 | 0.8066 | 0.5423 | 0.6014 | 0.5505 |

Finally, in the *randomly initialized specificity head* setting, 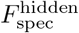is initialized from scratch rather than from the specificity-trained model, and is trained jointly with MLP_reg_. This setting measures the value of specificity pretraining for affinity prediction.

Table S8 shows that fine-tuning the pretrained specificity interaction head is critical for strong affinity prediction. Across all three test sets, the *unfrozen specificity head* variant achieves the best performance on both Spearman’s *ρ* and MAE, substantially outperforming the frozen variant. This confirms that the representation learned for specificity provides a strong initialization, but that additional task-specific adaptation is necessary because affinity regression must resolve finer distinctions among binding pairs than binary binder discrimination alone.

The *randomly initialized head* consistently underperforms the pretrained and fine-tuned variant, indicating that reusing specificity pretraining provides a clear advantage over learning the interaction representation from scratch. At the same time, the random-initialization variant is generally stronger than the fully frozen model, suggesting that affinity prediction does require optimization pressure on the interaction module itself rather than treating the specificity embedding as a fixed representation.

Adding ESM-based sequence fusion did not improve over the best *unfrozen specificity head* model on any test set. Although this variant remained competitive in rank correlation on AFF and HER2, it produced consistently higher MAE and markedly weaker performance on SKE. These results suggest that, after adaptation of the interaction module, the Protenix-derived structural representation was already sufficiently expressive for affinity prediction, with no additional benefit from sequence-level fusion in this setting.

### D.3 Developability Ablations

To understand how different information sources contribute to developability prediction, we perform a cumulative ablation study on the developability branch. The ablations are designed to progressively introduce (i) token-level information from the Protenix single representation, (ii) explicit pairwise structural information from the distogram output, (iii) lightweight task-specific adaptation of the shared Protenix structural representations, and (iv) additional sequence semantics from a frozen language model.

The first variant was a *single-only baseline*, which used only the pooled Protenix single representation,

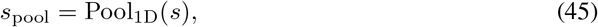

followed by a lightweight MLP regressor. This setting served as a sequence-dominant baseline within the Protenix feature space, as the single representation primarily captured token-level context.

Explicit pairwise structural information was then introduced through distogram-derived features from the Protenix distogram head. Let 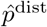 denote the distogram logits predicted from the pair representation. These logits were converted to probabilities and pooled into a pairwise feature *d*_pool_, giving the intermediate developability representation

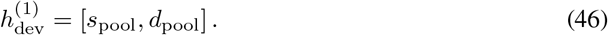

This variant, denoted *+ Distogram-based pair features*, augmented token-level features with explicit structural information.

**Table S9:** Cumulative ablation study of the developability branch. Each row adds one component on top of the previous setting. We report Spearman’s rank correlation coefficient (*ρ*) on four developability properties. Best and second-best results in each column are shown in **bold** and underlined, respectively.

| Ablation setting | Aggregation | Expression | Immunogenicity | Thermostability |
|---|---|---|---|---|
| Single-only baseline | 0.277 | 0.107 | 0.339 | 0.458 |
| + Distogram-based pair features | 0.338 | 0.217 | 0.469 | 0.621 |
| + Lightweight Pairformer adaptation | 0.318 | 0.285 | **0.493** | 0.720 |
| + ESM sequence fusion | **0.406** | **0.348** | 0.367 | **0.845** |

The third variant introduced a lightweight Pairformer refinement module to adapt the shared Protenix representations to developability prediction. Distogram-derived pairwise features were first injected into the pair representation,

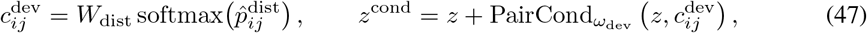

and then refine (*s, z*^cond^) with a single Pairformer block:

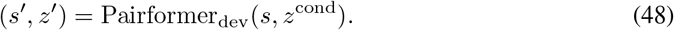

The refined representations are pooled as

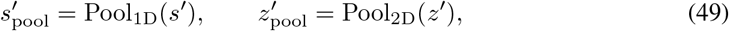

and used for scalar prediction. This setting was denoted *+ Lightweight Pairformer adaptation* and tested whether limited task-specific structural refinement improved developability prediction beyond fixed folding-derived features.

In the full model, the refined structural summaries were further fused with frozen ESM-2 embeddings of the heavy and light chains:

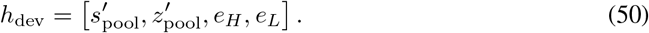

Within the full developability head 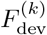, the resulting feature vector was mapped by the final MLP to a scalar prediction. This setting was denoted *+ ESM sequence fusion* and tested whether additional language-model sequence features improved antibody-centered property prediction beyond Protenix-derived structural features.

Table S9 shows that developability properties benefited from different representation components. Adding distogram-based pair features improved all four tasks, with the largest gain on immunogenicity, indicating that explicit pairwise structural information provided useful signals beyond token-level Protenix summaries.

*+ Lightweight Pairformer adaptation* further improved immunogenicity and thermostability, with immunogenicity reaching its best performance at this stage. This result suggested that immunogenicity benefited from task-specific refinement of the folding-derived representation, rather than from raw structural features alone.

*+ ESM sequence fusion* achieved the best results for aggregation, expression, and thermostability, improving Spearman’s *ρ* over the single-only baseline from 0.277 to 0.406, 0.107 to 0.348, and 0.458 to 0.845, respectively. These gains indicate that richer sequence-level semantics complemented Protenix-derived structural features for these intrinsic and manufacturability-related properties.

Overall, the ablation indicated that pairwise structural features improved most developability tasks, lightweight structural adaptation was particularly important for immunogenicity, and ESM sequence fusion provided the strongest overall gains across most endpoints.

## E Additional Results and Analysis

### E.1 Additional Comparisons with Protein Language Models

We extend the protein language model comparisons along two dimensions: parameter scale within the ESM-2 family [Lin et al., 2023] and encoder family through E1-600M [Jain et al., 2025] and ESMC-6B [Candido et al., 2026]. The scale experiments examine ESM models ranging from 8M to 3B parameters, while the additional encoder comparisons cover specificity, affinity, and developability prediction. For E1-600M and ESMC-6B, we follow the same fixed-length embedding pooling, lightweight prediction heads, task-specific training protocols, and validation-based checkpoint selection used for the language-model baselines.

**Table S10:**
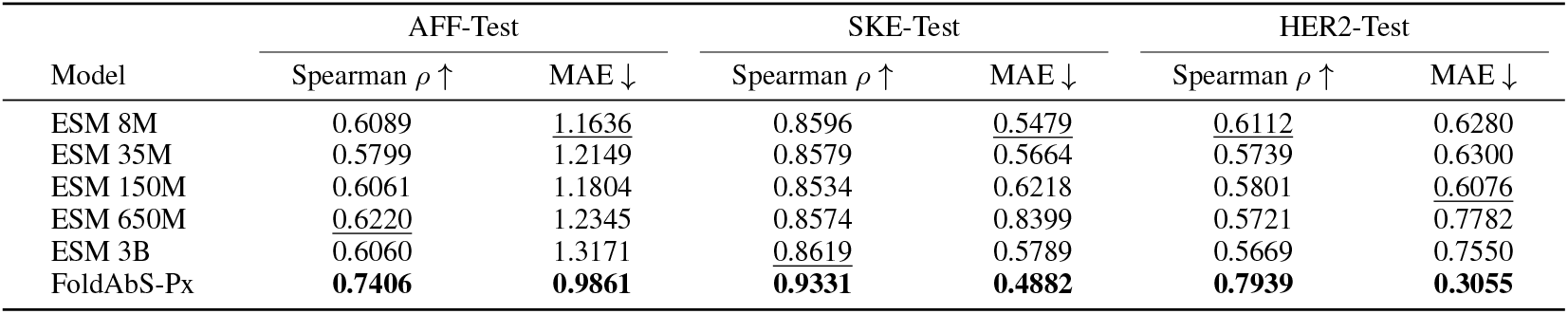
Effect of ESM model scale on affinity prediction. We compare FoldAbS-Px against ESM models of different sizes on three affinity benchmarks. Best and second-best results in each column are shown in **bold** and underlined, respectively.

| Model | AFF-Test |  | SKE-Test |  | HER2-Test |  |
|---|---|---|---|---|---|---|
|  | Spearman ρ ↑ | MAE ↓ | Spearman ρ ↑ | MAE ↓ | Spearman ρ ↑ | MAE ↓ |
| ESM 8M | 0.6089 | 1.1636 | 0.8596 | 0.5479 | 0.6112 | 0.6280 |
| ESM 35M | 0.5799 | 1.2149 | 0.8579 | 0.5664 | 0.5739 | 0.6300 |
| ESM 150M | 0.6061 | 1.1804 | 0.8534 | 0.6218 | 0.5801 | 0.6076 |
| ESM 650M | 0.6220 | 1.2345 | 0.8574 | 0.8399 | 0.5721 | 0.7782 |
| ESM 3B | 0.6060 | 1.3171 | 0.8619 | 0.5789 | 0.5669 | 0.7550 |
| FoldAbS-Px | **0.7406** | **0.9861** | **0.9331** | **0.4882** | **0.7939** | **0.3055** |

**Table S11:** Effect of ESM model scale on developability prediction. We report Spearman’s rank correlation coefficient (*ρ*) across four antibody properties. Best and second-best results in each column are shown in **bold** and underlined, respectively.

| Model | Aggregation | Expression | Immunogenicity | Thermostability |
|---|---|---|---|---|
| ESM 8M | 0.355 | 0.219 | 0.276 | 0.641 |
| ESM 35M | 0.353 | 0.116 | 0.304 | 0.765 |
| ESM 150M | 0.294 | 0.280 | 0.274 | 0.796 |
| ESM 650M | 0.362 | 0.311 | 0.202 | 0.784 |
| ESM 3B | 0.365 | 0.274 | 0.229 | 0.814 |
| FoldAbS-Px | **0.406** | **0.348** | **0.367** | **0.845** |

#### Scaling language models for affinity prediction

Table S10 compares FoldAbS-Px with ESM models of different scales on AFF-Test, SKE-Test, and HER2-Test. Across all three benchmarks, FoldAbS-Px substantially outperforms every ESM baseline in both Spearman’s *ρ* and MAE. On AFF-Test, the strongest ESM baseline reaches *ρ* = 0.6220, whereas FoldAbS-Px attains *ρ* = 0.7406 with a substantially lower MAE (0.9861 vs. 1.1636–1.3171). On SKE-Test, all ESM variants remain below the performance of FoldAbS-Px, with the best ESM model reaching *ρ* = 0.8619 compared with 0.9331 for FoldAbS-Px. On HER2-Test, the gap is even larger, where the best ESM model obtained *ρ* = 0.6112 while FoldAbS-Px achieved 0.7939. Notably, increasing ESM size does not produce monotonic gains: the 650M and 3B variants are not consistently better than the smaller models, and often yield worse MAE. These results indicate that the improvement of FoldAbS-Px cannot be attributed merely to model scale, but instead arises from reusing and adapting a structurally informed interaction representation.

#### Scaling language models for developability prediction

Table S11 reports the same analysis for developability prediction. Compared with ESM baselines, larger language models did not consistently improve performance across properties: ESM-650M performed slightly better on expression, ESM3B performed marginally better on aggregation and thermostability, and intermediate scales achieved the best results on some other properties. However, none of the ESM variants matched the overall performance of FoldAbS-Px. FoldAbS-Px achieved the best results on aggregation (0.406), expression (0.348), and thermostability (0.845), and outperformed all ESM models on immunogenicity (0.367 vs. 0.304 for the best ESM baseline). These results suggest that the main benefit of FoldAbS-Px did not arise from language-model scaling alone, but from combining Protenix-derived structural representations with lightweight task-specific adaptation.

**Table S12:**
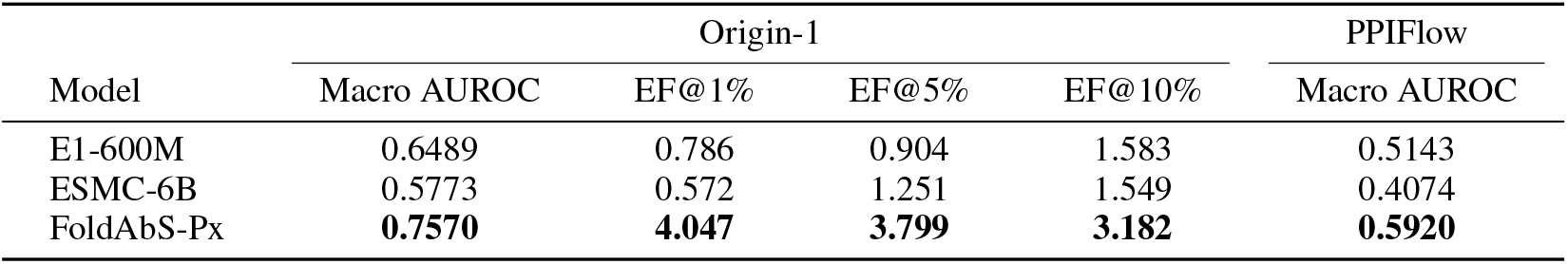
Specificity prediction with additional protein language models. We report macroaverage AUROC and enrichment factors on Origin-1, and macro-average AUROC on PPIFlow. Best results are shown in **bold**.

**Table S13:** Affinity prediction with additional protein language models. We report Spearman’s rank correlation coefficient (*ρ*) and MAE in the unified *pK*_*d*_ space. Best results are shown in **bold**.

|  | AFF-Test |  | SKE-Test |  | HER2-Test |  |
|---|---|---|---|---|---|---|
| Model | Spearman ρ ↑ | MAE ↓ | Spearman ρ ↑ | MAE ↓ | Spearman ρ ↑ | MAE ↓ |
| E1-600M | 0.6042 | 1.2227 | 0.8704 | 0.5519 | 0.6668 | 0.6860 |
| ESMC-6B | 0.6771 | 1.1686 | 0.8653 | 0.8179 | 0.6476 | 0.7895 |
| FoldAbS-Px | **0.7406** | **0.9861** | **0.9331** | **0.4882** | **0.7939** | **0.3055** |

#### Additional language models for specificity prediction

Table S12 compares FoldAbS-Px with E1-600M and ESMC-6B on Origin-1 and PPIFlow. FoldAbS-Px achieves the highest macro AUROC on both benchmarks, reaching 0.7570 on Origin-1 and 0.5920 on PPIFlow, compared with 0.6489 and 0.5143 for E1-600M, the stronger of the two additional encoders. FoldAbS-Px also achieves the highest early enrichment at all three Origin-1 screening rates. These results extend the specificity comparison across protein language model families and demonstrate a consistent advantage for task-adapted folding representations in binder discrimination and candidate prioritization.

#### Additional language models for affinity prediction

Table S13 reports affinity prediction results on AFF-Test, SKE-Test, and HER2-Test. FoldAbS-Px outperforms both additional encoders on all three benchmarks in both Spearman’s *ρ* and MAE. Compared with the stronger additional encoder on each benchmark, FoldAbS-Px increases Spearman’s *ρ* from 0.6771 to 0.7406 on AFF-Test, from 0.8704 to 0.9331 on SKE-Test, and from 0.6668 to 0.7939 on HER2-Test. The improvements in ranking accuracy are accompanied by lower absolute prediction errors across all three datasets, demonstrating the effectiveness of the adapted interaction representation for quantitative affinity prediction.

#### Additional language models for developability prediction

We compare E1-600M, ESMC-6B, and FoldAbS-Px using five-fold cross-validation with identical folds across methods. Table S14 reports the mean and standard deviation of Spearman’s *ρ* across folds. FoldAbS-Px achieves the highest mean correlation among these models on all four properties, covering aggregation, expression, immunogenicity, and thermostability. Together with the specificity and affinity results, these comparisons demonstrate that the performance advantage of FoldAbS-Px extends across protein language model families and across interaction-level and antibody-intrinsic prediction tasks.

### E.2 Specificity Prediction on the Cross-Antigen Benchmark

Specificity prediction was further evaluated on the challenging cross-antigen benchmark introduced by Smorodina et al. [2026]. This benchmark was derived from 106 experimentally annotated antibody– antigen pairs. Following the original protocol, antibodies and antigens were recombined by pair shuffling, with duplicate pairs removed, yielding 9,858 candidate pairs. Because several antigens were recognized by multiple antibodies, the final benchmark contained 110 positive pairs, resulting in severe class imbalance. Precision–recall AUC (PRAUC) was therefore used as the primary metric, following Smorodina et al. [2026], as it is more informative than AUROC in this setting.

Table S15 shows that FoldAbS-Px achieves the best PRAUC on this benchmark, outperforming both folding-model baselines and all language-model-based baselines. The improvement over Protenix and AF3 indicates that task-specific supervision remains beneficial even in a highly imbalanced recombined-pair setting. This behavior is further illustrated in Figure S1, where the score matrix produced by FoldAbS-Px exhibits substantially lower off-diagonal values than the native Protenix ipTM. FoldAbS-Px suppressed scores assigned to non-cognate antibody–antigen pairs more strongly, thereby reducing false positives in large candidate spaces. At the same time, the absolute PRAUC values remained low for all methods, including FoldAbS-Px, highlighting the difficulty of fine-grained antigen-specific recognition under extreme class imbalance. These results suggest that, although Protenix-based task adaptation improves screening quality, substantial room remains for future work on cross-antigen specificity prediction.

To directly assess antigen-conditioned discrimination, we hold each antibody fixed and compare its cognate and non-cognate antigen scores. For each cognate pair, the mean-decoy and hardest-decoy margins are defined as its score minus the mean and maximum scores over non-cognate antigens, respectively. As shown in Table S16(a), FoldAbS-Px increases the mean-decoy margin from 0.2029 to 0.2140 and reduces the score deficit relative to the hardest non-cognate antigen from 0.3047 to 0.0902.

We further control for native folding confidence by matching each cognate pair to the non-cognate pair for the same antibody with the closest Protenix ipTM. The median absolute ipTM difference between matched pairs is 0.0032. Under this matched comparison, FoldAbS-Px achieves antibodyand pair-level win rates of 62.3% and 62.7%, respectively, together with a positive mean margin of 0.0540 and median margin of 0.0115. The antibody-level bootstrap confidence intervals exclude 50% for both win rates and zero for both margins (Table S16(b)). These results demonstrate that FoldAbS-Px preferentially scores cognate interactions even when antibody identity and native folding confidence are controlled, establishing an antigen-conditioned screening signal beyond scalar ipTM.

### E.3 Early Enrichment on the PPIFlow Test Set

AUROC was used as the primary metric for PPIFlow because each target contained only 30 pairs, making low-rate enrichment sensitive to a few top-ranked examples. For completeness, enrichment factors at 5%, 10%, and 15% are reported in Table S17. The results were consistent with the main AUROC comparison: FoldAbS-Px matched or exceeded Protenix-v1 at lower screening rates, and both methods performed similarly at 15%.

### E.4 Effect of Pseudo-Negative Composition in Specificity Training

Specificity supervision used three pseudo-negative types: cross-antigen mismatches, CDRH3 mutation-based decoys, and natural CDRH3 swap-based decoys. To assess the effect of negative-pool composition on screening performance, the default FoldAbS-Px training recipe was compared with two variants: removal of natural CDRH3 swap-based negatives and an increased cross-antigen negative proportion from 0.3 to 0.5. All models were evaluated on Origin-1 using per-antigen AUROC, macro-average AUROC, and early enrichment factors.

Table S18 shows that the full FoldAbS-Px configuration yields the strongest overall specificity performance. In particular, it achieves the best macro AUROC and the best early enrichment at all screening rates, indicating that the full mixture of pseudo-negative types provides the most effective training signal for candidate prioritization. Removing natural CDRH3 swap-based negatives leads to a consistent drop in macro AUROC (from 0.757 to 0.739) and in all enrichment metrics, although it slightly improves performance on TNFRSF9. This suggests that natural CDRH3 swaps provide useful hard negatives that improve the model’s ability to suppress false positives beyond coarse antigen mismatch alone.

**Figure S1:**
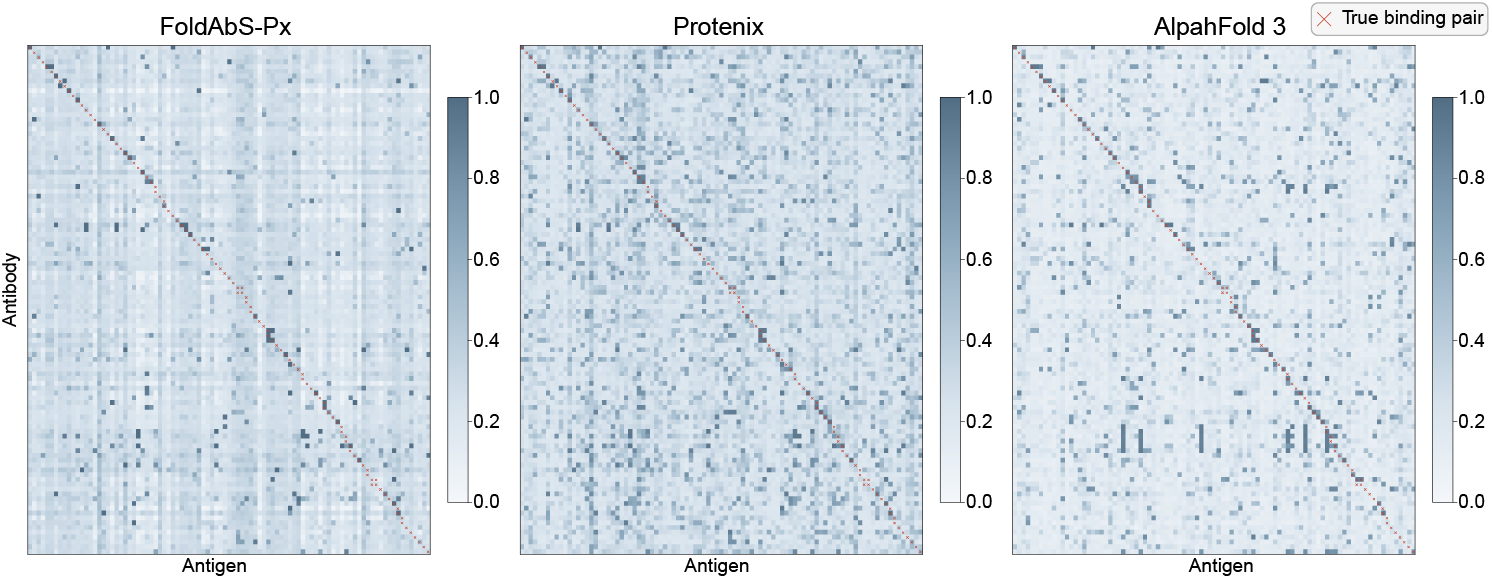
Score heatmaps on the cross-antigen benchmark. Rows and columns correspond to antibodies and antigens, respectively. Compared with the native Protenix ipTM, FoldAbS-Px assigns substantially lower scores to off-diagonal antibody–antigen pairs, indicating stronger suppression of non-cognate interactions and a lower false-positive tendency in the shuffled cross-antigen setting.

**Table S14:** Five-fold developability evaluation with additional protein language models. We report Spearman’s *ρ* as mean *±* standard deviation across identical folds. The highest mean for each property is shown in **bold**.

| Model | Aggregation | Expression | Immunogenicity | Thermostability |
|---|---|---|---|---|
| E1-600M | 0.2666 ± 0.1032 | 0.2440 ± 0.1046 | 0.3344 ± 0.0392 | 0.7403 ± 0.0576 |
| ESMC-6B | 0.1840 ± 0.0891 | 0.1346 ± 0.0883 | 0.3028 ± 0.1092 | 0.7282 ± 0.0550 |
| FoldAbS-Px | **0.3872** ± 0.0776 | **0.3458** ± 0.0681 | **0.3834** ± 0.1104 | **0.7876** ± 0.0710 |

**Table S15:** Specificity prediction on the cross-antigen benchmark of Smorodina et al. [2026]. We report PRAUC following the original evaluation protocol. Best and second-best results are shown in **bold** and underlined, respectively.

| Metric | FoldAbS-Px | Protenix | AF3 | ESM-650M | Ophiuchus-Ab | S3AI |
|---|---|---|---|---|---|---|
| PRAUC | **0.173** | <u>0.159</u> | 0.157 | 0.011 | 0.013 | 0.012 |

**Table S16:** Fixed-antibody and ipTM-matched analyses on the cross-antigen benchmark. (a) Cognate–decoy score margins with antibody identity held fixed. (b) Cognate preference after matching each cognate pair to a non-cognate pair for the same antibody with the closest Protenix ipTM. Confidence intervals are computed by antibody-level bootstrap resampling.

| *(a) Fixed-antibody score margins* |  |  |
| Metric | Protenix ipTM | FoldAbS-Px |
|---|---|---|
| Mean-decoy margin | 0.2029 | 0.2140 |
| Hardest-decoy margin | −0.3047 | −0.0902 |
| *(b) ipTM-matched cognate–decoy comparisons* |  |  |
| Metric | Estimate | 95% CI |
| Antibody win rate | 62.3% | [52.8%, 71.2%] |
| Pair win rate | 62.7% | [53.6%, 71.3%] |
| Mean margin | 0.0540 | [0.0218, 0.0870] |
| Median margin | 0.0115 | [0.0026, 0.0400] |

Increasing the proportion of cross-antigen negatives from 0.3 to 0.5 yields mixed behavior. While the larger ratio improves AUROC on a small number of antigen systems, including C5 and FXI, it substantially reduces the overall AUROC and degrades all enrichment metrics relative to the default setting. The drop is particularly pronounced on TSLP and IL36R, indicating that over-emphasizing coarse cross-antigen mismatches can weaken the model’s ability to distinguish more subtle nonbinding decoys. This interpretation is consistent with the training dynamics shown in Figure S2: the default ratio of 0.3 converges faster and more stably than the more aggressive 0.5 setting. Taken together, these results support the design choice of combining coarse cross-antigen negatives with harder CDRH3-based perturbations, rather than relying too heavily on cross-antigen mismatches alone.

### E.5 Per-Task versus Joint Training

Joint training was evaluated against the default per-task strategy. For affinity prediction, a single model was trained on the combined AFF, SKE, and HER2 datasets after *pK*_*d*_ standardization and evaluated separately on each benchmark. For developability prediction, the single Pairformer block was shared, while four property-specific MLP heads were used for aggregation, expression, immunogenicity, and thermostability.

**Table S17:**
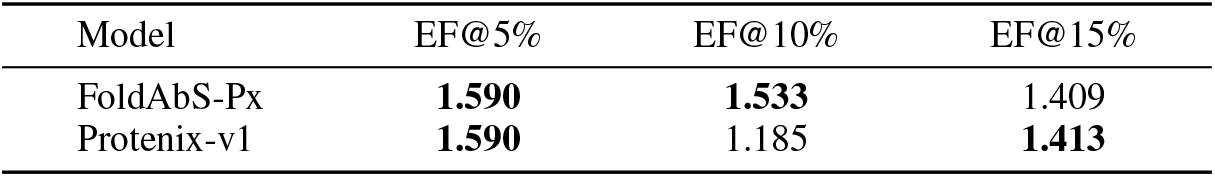
Early enrichment on the PPIFlow test set. We report enrichment factors at 5%, 10%, and 15% screening rates.

| Model | EF@5% | EF@10% | EF@15% |
|---|---|---|---|
| FoldAbS-Px | **1.590** | **1.533** | 1.409 |
| Protenix-v1 | **1.590** | 1.185 | **1.413** |

**Table S18:** Effect of pseudo-negative composition on specificity prediction. The default FoldAbS-Px setting uses a cross-antigen negative ratio of 0.3 together with mutation-based and natural CDRH3 swap-based negatives. “No natural CDRH3 swap” removes the swap-based decoy component, and “Higher cross-antigen ratio (0.5)” increases the proportion of cross-antigen mismatches during training. Best and second-best results in each column are shown in **bold** and underlined, respectively.

| Model | ACVR2B | C5 | FXI | HER2 | IL17A | IL36R | TNFRSF9 | TSLP | Macro Avg. | EF@1% | EF@5% | EF@10% |
|---|---|---|---|---|---|---|---|---|---|---|---|---|
| FoldAbS-Px (cross ratio 0.3) | **0.779** | 0.685 | 0.573 | **0.701** | **0.945** | **0.781** | 0.945 | **0.650** | **0.757** | **4.047** | **3.799** | **3.182** |
| No natural CDRH3 swap | 0.732 | 0.673 | 0.577 | 0.687 | 0.911 | 0.770 | **0.947** | 0.617 | 0.739 | 3.992 | 3.319 | 2.899 |
| Higher cross-antigen ratio (0.5) | 0.638 | **0.691** | **0.626** | 0.685 | 0.935 | 0.637 | 0.921 | 0.400 | 0.692 | 3.790 | 3.010 | 2.148 |

Tables S19 and S20 show that joint training did not consistently outperform task-specific training. For affinity prediction, per-dataset models achieved better Spearman’s *ρ* and MAE on all three benchmarks, indicating that *pK*_*d*_ standardization did not eliminate dataset heterogeneity across AFF, SKE, and HER2. Differences in sequence distribution, antigen composition, and supervision source, therefore, still favored dataset-specific adaptation.

For developability prediction, joint training improved aggregation from 0.406 to 0.440 but reduced performance on expression, immunogenicity, and thermostability. Thus, despite label normalization, developability properties remained heterogeneous and benefited from task-specific specialization. These results indicate that a shared numerical scale alone was insufficient to unify distinct antibody engineering tasks under a single training regime.

### E.6 Case Studies of Specificity Prediction

To qualitatively examine how the learned specificity signal of FoldAbS-Px differs from the native folding-model confidence signal, we visualize representative antibody–antigen pairs for which the two methods give markedly different scores. We focus on two complementary scenarios: (i) true binding pairs assigned high scores by FoldAbS-Px but relatively low iPTM by Protenix, and (ii) true non-binding pairs assigned low scores by FoldAbS-Px but high iPTM by Protenix. These examples illustrate how task-specific supervision reshapes the ranking signal beyond native structure-confidence estimates.

The contrast between Figure S3 and Figure S4 is consistent with the quantitative specificity results in the main text. On true binders, FoldAbS-Px remains highly confident even when native iPTM is only moderate, suggesting that folding-model confidence alone is not sufficient for binder ranking. On true non-binders, FoldAbS-Px systematically assigns much lower scores than Protenix, indicating stronger suppression of false positives. Taken together, these case studies support the view that task-specific supervision converts Protenix-derived structural signals into a screening metric that is more aligned with antibody specificity than native folding confidence alone.

### E.7 Specificity Prediction across Folding Backbones

To evaluate the transferability of FoldAbS across folding backbones, we instantiate the framework with Protenix-Mini [Gong et al., 2025] and Boltz2 [Passaro et al., 2025], denoted as FoldAbSPxMini and FoldAbS-Bz2, respectively. We retain the same task supervision, prediction-head design, and evaluation protocol, and compare each FoldAbS variant with its corresponding folding-model baseline. These experiments evaluate specificity prediction across Protenix model scales and across the Protenix and Boltz model families.

Table S21 shows that FoldAbS improves macro AUROC for both backbones on both benchmarks. With Protenix-Mini, FoldAbS increases macro AUROC from 0.6421 to 0.7305 on Origin-1 and from 0.5226 to 0.5753 on PPIFlow, corresponding to absolute gains of 0.0884 and 0.0527. The improvement extends to six of eight Origin-1 targets and four of seven PPIFlow targets, with one additional tie on PPIFlow. With Boltz2, FoldAbS increases macro AUROC from 0.6738 to 0.7333 on Origin-1 and from 0.5225 to 0.5896 on PPIFlow, yielding absolute gains of 0.0595 and 0.0671. FoldAbS-Bz2 improves performance on seven of eight Origin-1 targets and six of seven PPIFlow targets, demonstrating broad gains across antigen systems.

**Figure S2:**
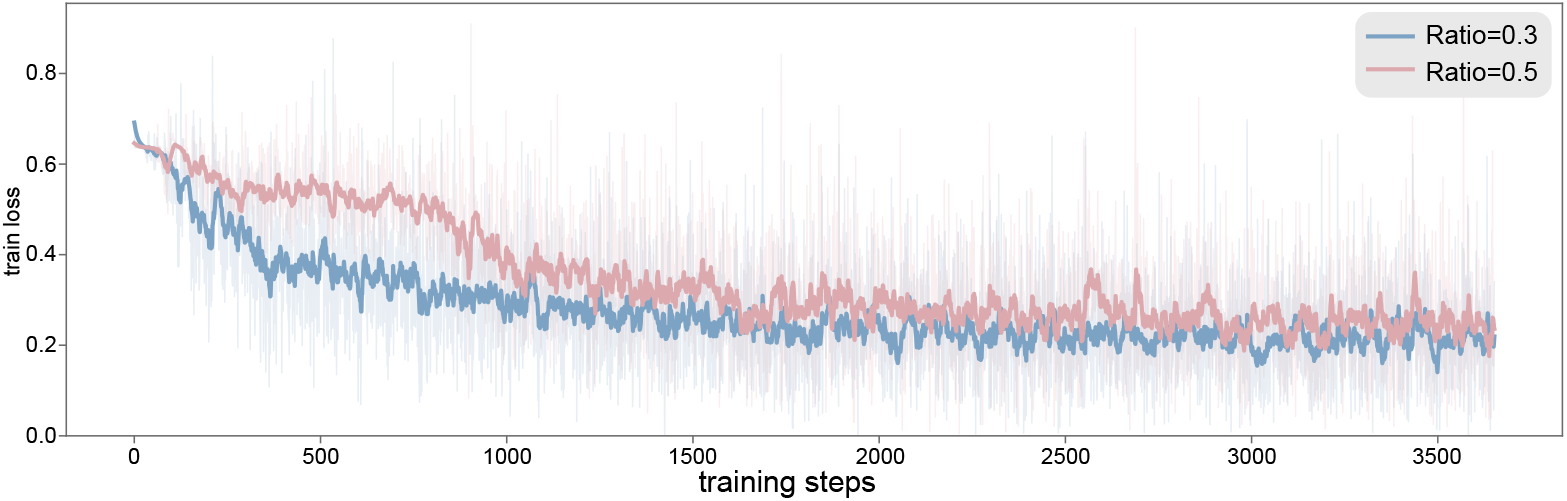
Training loss curves under different cross-antigen negative ratios. We compare the default specificity training configuration using a cross-antigen negative ratio of 0.3 with a more aggressive setting using a ratio of 0.5. The 0.3 setting converges faster and more stably, supporting the final evaluation results in Table S18.

**Table S19:** Affinity prediction under per-dataset versus joint training. All datasets are mapped to the shared *pK*_*d*_ space, while evaluation is performed separately on each test set. Best results in each column are shown in **bold**.

|  | AFF-Test |  | SKE-Test |  | HER2-Test |  |
|---|---|---|---|---|---|---|
| Training setting | Spearman ρ ↑ | MAE ↓ | Spearman ρ ↑ | MAE ↓ | Spearman ρ ↑ | MAE ↓ |
| FoldAbS-Px (per-dataset) | **0.7406** | **0.9861** | **0.9331** | **0.4882** | **0.7939** | **0.3055** |
| FoldAbS-Px (joint) | 0.7169 | 1.0231 | 0.8024 | 0.7399 | 0.7308 | 0.3787 |

The gains also translate into improved early enrichment. As shown in Table S22, FoldAbS-Bz2 increases EF@1%, EF@5%, and EF@10% from 2.9890, 2.6251, and 2.1431 to 4.1261, 3.4383, and 2.7797, respectively, improving binder recovery across all three screening rates. Together with the Protenix-v1 results in the main text, these experiments demonstrate that supervised adaptation of folding-derived representations consistently improves specificity screening across model scales and folding-model families.

### E.8 Five-Fold Cross-Validation for Developability Prediction

We evaluate FoldAbS-Px and all developability baselines using five-fold cross-validation with identical folds across methods. The comparison includes ESM-650M [Lin et al., 2023], E1-600M [Jain et al., 2025], ESMC-6B [Candido et al., 2026], AntiBERTy [Ruffolo et al., 2021], Ophiuchus-Ab [Zhu et al., 2026], and three IgFold variants [Ruffolo et al., 2023]. Table S23 reports Spearman’s rank correlation coefficient (*ρ*) as mean *±* standard deviation across folds.

FoldAbS-Px achieves the highest reported mean correlation on all four properties. Its largest advantage over the strongest competing mean is observed on aggregation, where it reaches 0.3872 compared with 0.3377 for Ophiuchus-Ab. For expression, FoldAbS-Px achieves 0.3458 compared with 0.3309 for IgFold bert; for thermostability, it reaches 0.7876 compared with 0.7641 for Ophiuchus-Ab. On immunogenicity, FoldAbS-Px achieves 0.3834, closely matching AntiBERTy (0.383) and OphiuchusAb (0.3816). The five-fold results therefore reproduce the overall advantage observed in the original split on aggregation, expression, and thermostability, while identifying comparable immunogenicity performance among the leading models. These results demonstrate that the effectiveness of FoldAbSPx extends across data partitions and heterogeneous developability properties.

**Table S20:** Developability prediction under per-task versus joint training. All properties are normalized to a shared zero-mean unit-variance scale during training. Best results in each column are shown in **bold**.

| Training setting | Aggregation | Expression | Immunogenicity | Thermostability |
|---|---|---|---|---|
| FoldAbS-Px (per-task) | 0.406 | **0.348** | **0.367** | **0.845** |
| FoldAbS-Px (joint) | **0.440** | 0.329 | 0.264 | 0.822 |

**Figure S3:**
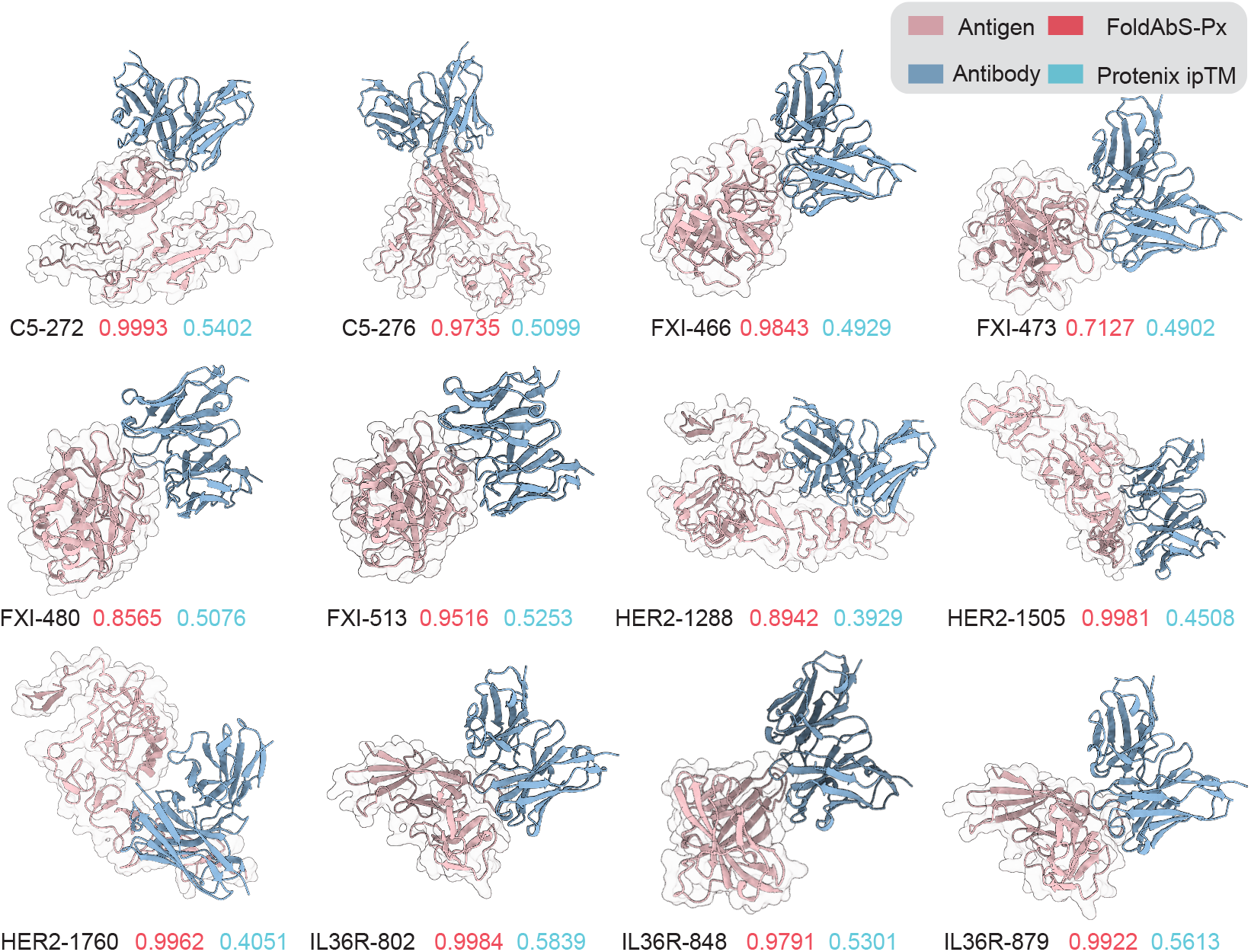
Binding case studies with high FoldAbS-Px scores but relatively low Protenix ipTM. Each example corresponds to a true binding antibody–antigen pair. Across these cases, FoldAbS-Px consistently assigns high specificity scores, whereas the native Protenix ipTM remains only moderate. This pattern suggests that the learned specificity head can recover true binders that are not strongly favored by native folding-model confidence alone.

## F Broader Impacts

This work presents a unified antibody-screening framework based on protein folding models for specificity, affinity, and developability prediction. The framework may support more efficient therapeutic antibody discovery by improving computational candidate prioritization. No substantial negative societal impacts were identified, although computational predictions should not replace experimental validation and should be interpreted with appropriate caution.

**Figure S4:**
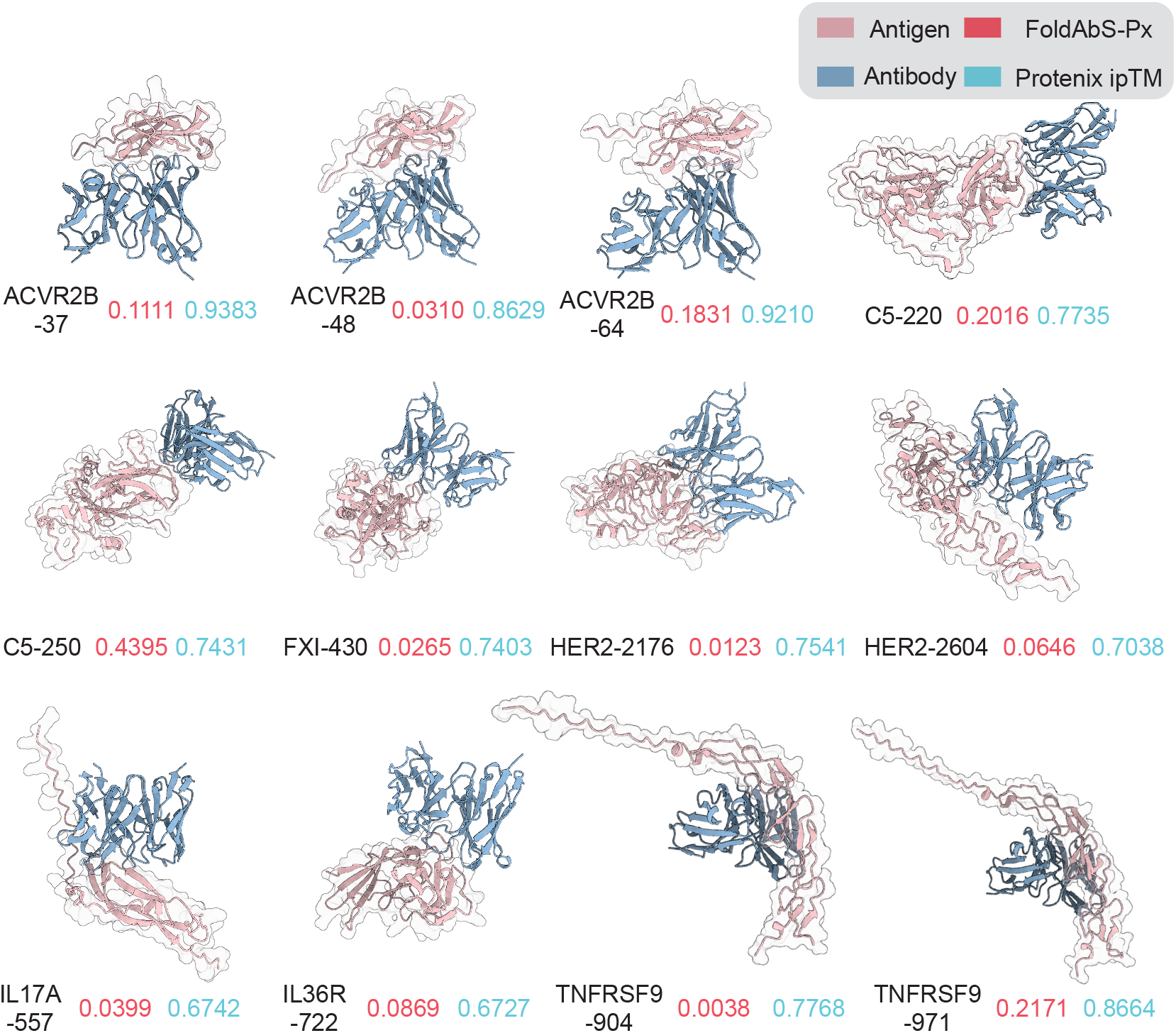
Non-binding case studies with low FoldAbS-Px scores but high Protenix ipTM. Each example corresponds to a true non-binding antibody–antigen pair. In these cases, Protenix assigns relatively high iPTM values, whereas FoldAbS-Px suppresses the scores substantially. This behavior indicates that the learned specificity head is better calibrated for rejecting non-cognate pairs, thereby reducing false positives in screening.

**Table S21:** Specificity prediction across folding backbones. We report per-target and macro-average AUROC on (a) Origin-1 and (b) PPIFlow. Bold values indicate the better result within each matched backbone pair; ties are highlighted for both methods.

| *(a) Origin-1* |  |  |  |  |  |  |  |  |  |
| Model | ACVR2B | C5 | FXI | HER2 | IL17A | IL36R | TNFRSF9 | TSLP | Macro Avg. |
|---|---|---|---|---|---|---|---|---|---|
| Protenix-Mini | 0.5429 | **0.6931** | 0.3759 | 0.4465 | 0.8092 | **0.7349** | 0.9145 | 0.6197 | 0.6421 |
| FoldAbS-PxMini | **0.6210** | 0.6214 | **0.6851** | **0.6765** | **0.9037** | 0.6498 | **0.9397** | **0.7471** | **0.7305** |
| Boltz2 | 0.6369 | 0.7663 | 0.5958 | 0.5557 | **0.7861** | 0.7234 | 0.7940 | 0.5321 | 0.6738 |
| FoldAbS-Bz2 | **0.6697** | **0.8069** | **0.6040** | **0.6413** | 0.7534 | **0.8062** | **0.8340** | **0.7507** | **0.7333** |
| *(b) PPIFlow* |  |  |  |  |  |  |  |  |  |
| Model | 1433E | BHRF1 | CCL2 | EFNA1 | HNMT | PDGFR | S100A4 | Macro Avg. |  |
| Protenix-Mini | 0.5208 | **0.7760** | 0.2986 | 0.5291 | **0.3819** | **0.5920** | 0.5600 | 0.5226 |  |
| FoldAbS-PxMini | **0.7431** | 0.7520 | **0.4299** | **0.6243** | **0.3819** | 0.4880 | **0.6080** | **0.5753** |  |
| Boltz2 | **0.6806** | 0.5600 | 0.3756 | 0.6825 | 0.3750 | 0.4880 | 0.4960 | 0.5225 |  |
| FoldAbS-Bz2 | 0.5486 | **0.6880** | **0.4434** | **0.6931** | **0.5625** | **0.5280** | **0.6640** | **0.5896** |  |

**Table S22:** Early enrichment with the Boltz2 backbone on Origin-1. We report macro-average enrichment factors at 1%, 5%, and 10% screening rates. Best results are shown in **bold**.

| Model | EF@1% | EF@5% | EF@10% |
|---|---|---|---|
| Boltz2 | 2.9890 | 2.6251 | 2.1431 |
| FoldAbS-Bz2 | **4.1261** | **3.4383** | **2.7797** |

**Table S23:** Five-fold cross-validation for developability prediction. We report Spearman’s *ρ* as mean *±* standard deviation across identical folds for all methods. The highest reported mean for each property is shown in **bold**.

| Model | Aggregation | Expression | Immunogenicity | Thermostability |
|---|---|---|---|---|
| FoldAbS-Px | **0.3872** ± 0.0776 | **0.3458** ± 0.0681 | **0.3834** ± 0.1104 | **0.7876** ± 0.0710 |
| ESM-650M | 0.3320 ± 0.0738 | 0.3133 ± 0.0930 | 0.3252 ± 0.1076 | 0.7094 ± 0.0553 |
| E1-600M | 0.2666 ± 0.1032 | 0.2440 ± 0.1046 | 0.3344 ± 0.0392 | 0.7403 ± 0.0576 |
| ESMC-6B | 0.1840 ± 0.0891 | 0.1346 ± 0.0883 | 0.3028 ± 0.1092 | 0.7282 ± 0.0550 |
| AntiBERTy | 0.184 ± 0.125 | 0.309 ± 0.136 | 0.383 ± 0.155 | 0.696 ± 0.072 |
| Ophiuchus-Ab | 0.3377 ± 0.1167 | 0.3129 ± 0.1168 | 0.3816 ± 0.1792 | 0.7641 ± 0.0596 |
| IgFold bert | 0.2248 ± 0.0401 | 0.3309 ± 0.1324 | 0.3649 ± 0.2033 | 0.6884 ± 0.0757 |
| IgFold struc | 0.105 ± 0.081 | 0.156 ± 0.089 | 0.211 ± 0.023 | 0.491 ± 0.042 |
| IgFold gt | 0.179 ± 0.112 | 0.155 ± 0.051 | 0.225 ± 0.150 | 0.489 ± 0.051 |

## NeurIPS Paper Checklist

The checklist is designed to encourage best practices for responsible machine learning research, addressing issues of reproducibility, transparency, research ethics, and societal impact. Do not remove the checklist: **The papers not including the checklist will be desk rejected**. The checklist should follow the references and follow the (optional) supplemental material. The checklist does NOT count towards the page limit.

Please read the checklist guidelines carefully for information on how to answer these questions. For each question in the checklist:

- You should answer [Yes], [No], or [N/A].
- [N/A] means either that the question is Not Applicable for that particular paper or the relevant information is Not Available.
- Please provide a short (1–2 sentence) justification right after your answer (even for [N/A]).

### The checklist answers are an integral part of your paper submission

They are visible to the reviewers, area chairs, senior area chairs, and ethics reviewers. You will also be asked to include it (after eventual revisions) with the final version of your paper, and its final version will be published with the paper.

The reviewers of your paper will be asked to use the checklist as one of the factors in their evaluation. While [Yes] is generally preferable to [No], it is perfectly acceptable to answer [No] provided a proper justification is given (e.g., error bars are not reported because it would be too computationally expensive” or “we were unable to find the license for the dataset we used”). In general, answering [No] or [N/A] is not grounds for rejection. While the questions are phrased in a binary way, we acknowledge that the true answer is often more nuanced, so please just use your best judgment and write a justification to elaborate. All supporting evidence can appear either in the main paper or the supplemental material, provided in appendix. If you answer [Yes] to a question, in the justification please point to the section(s) where related material for the question can be found.

IMPORTANT, please:

- **Delete this instruction block, but keep the section heading “NeurIPS Paper Checklist”**,
- **Keep the checklist subsection headings, questions/answers and guidelines below**.
- **Do not modify the questions and only use the provided macros for your answers**.

#### 1. Claims

Question: Do the main claims made in the abstract and introduction accurately reflect the paper’s contributions and scope?

Answer: [Yes]

Justification: We have made clear our paper’s research focus and contributions, with key contributions listed at the end of the introduction.

Guidelines:

- The answer [N/A] means that the abstract and introduction do not include the claims made in the paper.
- The abstract and/or introduction should clearly state the claims made, including the contributions made in the paper and important assumptions and limitations. A [No] or [N/A] answer to this question will not be perceived well by the reviewers.
- The claims made should match theoretical and experimental results, and reflect how much the results can be expected to generalize to other settings.
- It is fine to include aspirational goals as motivation as long as it is clear that these goals are not attained by the paper.

#### 2. Limitations

Question: Does the paper discuss the limitations of the work performed by the authors? Answer: [Yes]

Justification: We have discussed the limitations in Section 5 Guidelines:

- The answer [N/A] means that the paper has no limitation while the answer [No] means that the paper has limitations, but those are not discussed in the paper.
- The authors are encouraged to create a separate “Limitations” section in their paper.
- The paper should point out any strong assumptions and how robust the results are to violations of these assumptions (e.g., independence assumptions, noiseless settings, model well-specification, asymptotic approximations only holding locally). The authors should reflect on how these assumptions might be violated in practice and what the implications would be.
- The authors should reflect on the scope of the claims made, e.g., if the approach was only tested on a few datasets or with a few runs. In general, empirical results often depend on implicit assumptions, which should be articulated.
- The authors should reflect on the factors that influence the performance of the approach. For example, a facial recognition algorithm may perform poorly when image resolution is low or images are taken in low lighting. Or a speech-to-text system might not be used reliably to provide closed captions for online lectures because it fails to handle technical jargon.
- The authors should discuss the computational efficiency of the proposed algorithms and how they scale with dataset size.
- If applicable, the authors should discuss possible limitations of their approach to address problems of privacy and fairness.
- While the authors might fear that complete honesty about limitations might be used by reviewers as grounds for rejection, a worse outcome might be that reviewers discover limitations that aren’t acknowledged in the paper. The authors should use their best judgment and recognize that individual actions in favor of transparency play an important role in developing norms that preserve the integrity of the community. Reviewers will be specifically instructed to not penalize honesty concerning limitations.

#### 3. Theory assumptions and proofs

Question: For each theoretical result, does the paper provide the full set of assumptions and a complete (and correct) proof?

Answer: [N/A]

Justification: This paper does not propose any theorem. Therefore, it does not include any proof.

Guidelines:

- The answer [N/A] means that the paper does not include theoretical results.
- All the theorems, formulas, and proofs in the paper should be numbered and crossreferenced.
- All assumptions should be clearly stated or referenced in the statement of any theorems.
- The proofs can either appear in the main paper or the supplemental material, but if they appear in the supplemental material, the authors are encouraged to provide a short proof sketch to provide intuition.
- Inversely, any informal proof provided in the core of the paper should be complemented by formal proofs provided in appendix or supplemental material.
- Theorems and Lemmas that the proof relies upon should be properly referenced.

#### 4. Experimental result reproducibility

Question: Does the paper fully disclose all the information needed to reproduce the main experimental results of the paper to the extent that it affects the main claims and/or conclusions of the paper (regardless of whether the code and data are provided or not)?

Answer: [Yes]

Justification: We have provided full data processing details in Appendix B and presented the implementation of baseline methods and our approach in Appendix A.

Guidelines:

- The answer [N/A] means that the paper does not include experiments.
- If the paper includes experiments, a [No] answer to this question will not be perceived well by the reviewers: Making the paper reproducible is important, regardless of whether the code and data are provided or not.
- If the contribution is a dataset and/or model, the authors should describe the steps taken to make their results reproducible or verifiable.
- Depending on the contribution, reproducibility can be accomplished in various ways. For example, if the contribution is a novel architecture, describing the architecture fully might suffice, or if the contribution is a specific model and empirical evaluation, it may be necessary to either make it possible for others to replicate the model with the same dataset, or provide access to the model. In general. releasing code and data is often one good way to accomplish this, but reproducibility can also be provided via detailed instructions for how to replicate the results, access to a hosted model (e.g., in the case of a large language model), releasing of a model checkpoint, or other means that are appropriate to the research performed.
- While NeurIPS does not require releasing code, the conference does require all submissions to provide some reasonable avenue for reproducibility, which may depend on the nature of the contribution. For example
  a. If the contribution is primarily a new algorithm, the paper should make it clear how to reproduce that algorithm.
  b. If the contribution is primarily a new model architecture, the paper should describe the architecture clearly and fully.
  c. If the contribution is a new model (e.g., a large language model), then there should either be a way to access this model for reproducing the results or a way to reproduce the model (e.g., with an open-source dataset or instructions for how to construct the dataset).
  d. We recognize that reproducibility may be tricky in some cases, in which case authors are welcome to describe the particular way they provide for reproducibility. In the case of closed-source models, it may be that access to the model is limited in some way (e.g., to registered users), but it should be possible for other researchers to have some path to reproducing or verifying the results.

#### 5. Open access to data and code

Question: Does the paper provide open access to the data and code, with sufficient instructions to faithfully reproduce the main experimental results, as described in supplemental material?

Answer: [No]

Justification: The source code will be made publicly available upon acceptance. Guidelines:

- The answer [N/A] means that paper does not include experiments requiring code.
- Please see the NeurIPS code and data submission guidelines (https://neurips.cc/public/guides/CodeSubmissionPolicy) for more details.
- While we encourage the release of code and data, we understand that this might not be possible, so [No] is an acceptable answer. Papers cannot be rejected simply for not including code, unless this is central to the contribution (e.g., for a new open-source benchmark).
- The instructions should contain the exact command and environment needed to run to reproduce the results. See the NeurIPS code and data submission guidelines (https://neurips.cc/public/guides/CodeSubmissionPolicy) for more details.
- The authors should provide instructions on data access and preparation, including how to access the raw data, preprocessed data, intermediate data, and generated data, etc.
- The authors should provide scripts to reproduce all experimental results for the new proposed method and baselines. If only a subset of experiments are reproducible, they should state which ones are omitted from the script and why.
- At submission time, to preserve anonymity, the authors should release anonymized versions (if applicable).
- Providing as much information as possible in supplemental material (appended to the paper) is recommended, but including URLs to data and code is permitted.

#### 6. Experimental setting/details

Question: Does the paper specify all the training and test details (e.g., data splits, hyperparameters, how they were chosen, type of optimizer) necessary to understand the results?

Answer: [Yes]

Justification: We have provided full data processing details in Appendix B and presented the implementation of baseline methods and our approach in Appendix A.

Guidelines:

- The answer [N/A] means that the paper does not include experiments.
- The experimental setting should be presented in the core of the paper to a level of detail that is necessary to appreciate the results and make sense of them.
- The full details can be provided either with the code, in appendix, or as supplemental material.

#### 7. Experiment statistical significance

Question: Does the paper report error bars suitably and correctly defined or other appropriate information about the statistical significance of the experiments?

Answer: [No]

Justification: We follow the evaluation protocol of previous works, which does not involve reporting the error bars.

Guidelines:

- The answer [N/A] means that the paper does not include experiments.
- The authors should answer [Yes] if the results are accompanied by error bars, confidence intervals, or statistical significance tests, at least for the experiments that support the main claims of the paper.
- The factors of variability that the error bars are capturing should be clearly stated (for example, train/test split, initialization, random drawing of some parameter, or overall run with given experimental conditions).
- The method for calculating the error bars should be explained (closed form formula, call to a library function, bootstrap, etc.)
- The assumptions made should be given (e.g., Normally distributed errors).
- It should be clear whether the error bar is the standard deviation or the standard error of the mean.
- It is OK to report 1-sigma error bars, but one should state it. The authors should preferably report a 2-sigma error bar than state that they have a 96% CI, if the hypothesis of Normality of errors is not verified.
- For asymmetric distributions, the authors should be careful not to show in tables or figures symmetric error bars that would yield results that are out of range (e.g., negative error rates).
- If error bars are reported in tables or plots, the authors should explain in the text how they were calculated and reference the corresponding figures or tables in the text.

#### 8. Experiments compute resources

Question: For each experiment, does the paper provide sufficient information on the computer resources (type of compute workers, memory, time of execution) needed to reproduce the experiments?

Answer: [Yes]

Justification: We have provided sufficient information on the computer resources in Appendix A.

Guidelines:

- The answer [N/A] means that the paper does not include experiments.
- The paper should indicate the type of compute workers CPU or GPU, internal cluster, or cloud provider, including relevant memory and storage.
- The paper should provide the amount of compute required for each of the individual experimental runs as well as estimate the total compute.
- The paper should disclose whether the full research project required more compute than the experiments reported in the paper (e.g., preliminary or failed experiments that didn’t make it into the paper).

#### 9. Code of ethics

Question: Does the research conducted in the paper conform, in every respect, with the NeurIPS Code of Ethics https://neurips.cc/public/EthicsGuidelines?

Answer: [Yes]

Justification: The research conducted in the paper conform, in every respect, with the NeurIPS Code of Ethics.

Guidelines:

- The answer [N/A] means that the authors have not reviewed the NeurIPS Code of Ethics.
- If the authors answer [No], they should explain the special circumstances that require a deviation from the Code of Ethics.
- The authors should make sure to preserve anonymity (e.g., if there is a special consideration due to laws or regulations in their jurisdiction).

#### 10. Broader impacts

Question: Does the paper discuss both potential positive societal impacts and negative societal impacts of the work performed?

Answer: [Yes]

Justification: We have discussed the broader impacts in Appendix F Guidelines:

- The answer [N/A] means that there is no societal impact of the work performed.
- If the authors answer [N/A] or [No], they should explain why their work has no societal impact or why the paper does not address societal impact.
- Examples of negative societal impacts include potential malicious or unintended uses (e.g., disinformation, generating fake profiles, surveillance), fairness considerations (e.g., deployment of technologies that could make decisions that unfairly impact specific groups), privacy considerations, and security considerations.
- The conference expects that many papers will be foundational research and not tied to particular applications, let alone deployments. However, if there is a direct path to any negative applications, the authors should point it out. For example, it is legitimate to point out that an improvement in the quality of generative models could be used to generate Deepfakes for disinformation. On the other hand, it is not needed to point out that a generic algorithm for optimizing neural networks could enable people to train models that generate Deepfakes faster.
- The authors should consider possible harms that could arise when the technology is being used as intended and functioning correctly, harms that could arise when the technology is being used as intended but gives incorrect results, and harms following from (intentional or unintentional) misuse of the technology.
- If there are negative societal impacts, the authors could also discuss possible mitigation strategies (e.g., gated release of models, providing defenses in addition to attacks, mechanisms for monitoring misuse, mechanisms to monitor how a system learns from feedback over time, improving the efficiency and accessibility of ML).

#### 11. Safeguards

Question: Does the paper describe safeguards that have been put in place for responsible release of data or models that have a high risk for misuse (e.g., pre-trained language models, image generators, or scraped datasets)?

Answer: [N/A]

Justification: The paper poses no such risks. Guidelines:

- The answer [N/A] means that the paper poses no such risks.
- Released models that have a high risk for misuse or dual-use should be released with necessary safeguards to allow for controlled use of the model, for example by requiring that users adhere to usage guidelines or restrictions to access the model or implementing safety filters.
- Datasets that have been scraped from the Internet could pose safety risks. The authors should describe how they avoided releasing unsafe images.
- We recognize that providing effective safeguards is challenging, and many papers do not require this, but we encourage authors to take this into account and make a best faith effort.

#### 12. Licenses for existing assets

Question: Are the creators or original owners of assets (e.g., code, data, models), used in the paper, properly credited and are the license and terms of use explicitly mentioned and properly respected?

Answer: [Yes]

Justification: The original papers that introduce models and datasets used in the paper are already properly cited.

Guidelines:

- The answer [N/A] means that the paper does not use existing assets.
- The authors should cite the original paper that produced the code package or dataset.
- The authors should state which version of the asset is used and, if possible, include a URL.
- The name of the license (e.g., CC-BY 4.0) should be included for each asset.
- For scraped data from a particular source (e.g., website), the copyright and terms of service of that source should be provided.
- If assets are released, the license, copyright information, and terms of use in the package should be provided. For popular datasets, paperswithcode.com/datasets has curated licenses for some datasets. Their licensing guide can help determine the license of a dataset.
- For existing datasets that are re-packaged, both the original license and the license of the derived asset (if it has changed) should be provided.
- If this information is not available online, the authors are encouraged to reach out to the asset’s creators.

#### 13. New assets

Question: Are new assets introduced in the paper well documented and is the documentation provided alongside the assets?

Answer: [N/A]

Justification: Currently, the paper does not release new assets. Our source code will be released upon the acceptance of the paper with comprehensive documents.

Guidelines:

- The answer [N/A] means that the paper does not release new assets.
- Researchers should communicate the details of the dataset/code/model as part of their submissions via structured templates. This includes details about training, license, limitations, etc.
- The paper should discuss whether and how consent was obtained from people whose asset is used.
- At submission time, remember to anonymize your assets (if applicable). You can either create an anonymized URL or include an anonymized zip file.

#### 14. Crowdsourcing and research with human subjects

Question: For crowdsourcing experiments and research with human subjects, does the paper include the full text of instructions given to participants and screenshots, if applicable, as well as details about compensation (if any)?

Answer: [N/A]

Justification: The paper does not involve crowdsourcing nor research with human subjects. Guidelines:

- The answer [N/A] means that the paper does not involve crowdsourcing nor research with human subjects.
- Including this information in the supplemental material is fine, but if the main contribution of the paper involves human subjects, then as much detail as possible should be included in the main paper.
- According to the NeurIPS Code of Ethics, workers involved in data collection, curation, or other labor should be paid at least the minimum wage in the country of the data collector.

#### 15. Institutional review board (IRB) approvals or equivalent for research with human subjects

Question: Does the paper describe potential risks incurred by study participants, whether such risks were disclosed to the subjects, and whether Institutional Review Board (IRB) approvals (or an equivalent approval/review based on the requirements of your country or institution) were obtained?

Answer: [N/A]

Justification: This paper does not incur such risks. Guidelines:

- The answer [N/A] means that the paper does not involve crowdsourcing nor research with human subjects.
- Depending on the country in which research is conducted, IRB approval (or equivalent) may be required for any human subjects research. If you obtained IRB approval, you should clearly state this in the paper.
- We recognize that the procedures for this may vary significantly between institutions and locations, and we expect authors to adhere to the NeurIPS Code of Ethics and the guidelines for their institution.
- For initial submissions, do not include any information that would break anonymity (if applicable), such as the institution conducting the review.

#### 16. Declaration of LLM usage

Question: Does the paper describe the usage of LLMs if it is an important, original, or non-standard component of the core methods in this research? Note that if the LLM is used only for writing, editing, or formatting purposes and does *not* impact the core methodology, scientific rigor, or originality of the research, declaration is not required.

Answer: [N/A]

Justification: The core method development in this research does not involve LLMs as any important, original, or non-standard components.

Guidelines:

- The answer [N/A] means that the core method development in this research does not involve LLMs as any important, original, or non-standard components.
- Please refer to our LLM policy in the NeurIPS handbook for what should or should not be described.

## Notes

### Competing Interest Statement

The authors have declared no competing interest.

